# Profile-likelihood Bayesian model averaging for two-sample summary data Mendelian randomization in the presence of horizontal pleiotropy

**DOI:** 10.1101/2020.02.11.943712

**Authors:** Chin Yang Shapland, Qingyuan Zhao, Jack Bowden

## Abstract

Two-sample summary data Mendelian randomisation is a popular method for assessing causality in epidemiology, by using genetic variants as instrumental variables. If genes exert pleiotropic effects on the outcome not entirely through the exposure of interest, this can lead to heterogeneous and (potentially) biased estimates of causal effect. We investigate the use of Bayesian model averaging to preferentially search the space of models with the highest posterior likelihood. We develop a Metropolis-Hasting algorithm to perform the search using the recently developed Robust Adjusted Profile Likelihood of Zhao et al as the basis for defining a posterior distribution that efficiently accounts for pleiotropic and weak instrument bias. We demonstrate how our general modelling approach can be extended from a standard one-parameter causal model to a two-parameter model, which allows a large proportion of SNPs to violate the Instrument Strength Independent of Direct Effect assumption. We use Monte Carlo simulations to illustrate our methods and compare it to several related approaches. We finish by applying our approach in practice to investigate the causal role of cholesterol on the development age-related macular degeneration.

## 1 Introduction

The capacity of traditional observational epidemiology to reliably infer whether a health exposure causally influences a disease rests on its ability to appropriately measure and adjust for factors which jointly predict (or confound) the exposure-outcome relationship. Mendelian randomization (MR) [1] avoids bias from unmeasured confounding by using genetic variants as instrumental variables (IVs) [2]. For the approach to be valid for testing causality, each specific IV must be robustly associated with the exposure (assumption IV1), independent of any confounders of the exposure and outcome (IV2) and be independent of the outcome given the exposure and the confounders (IV3), as illustrated by Figure 1a.

**Fig. 1:**
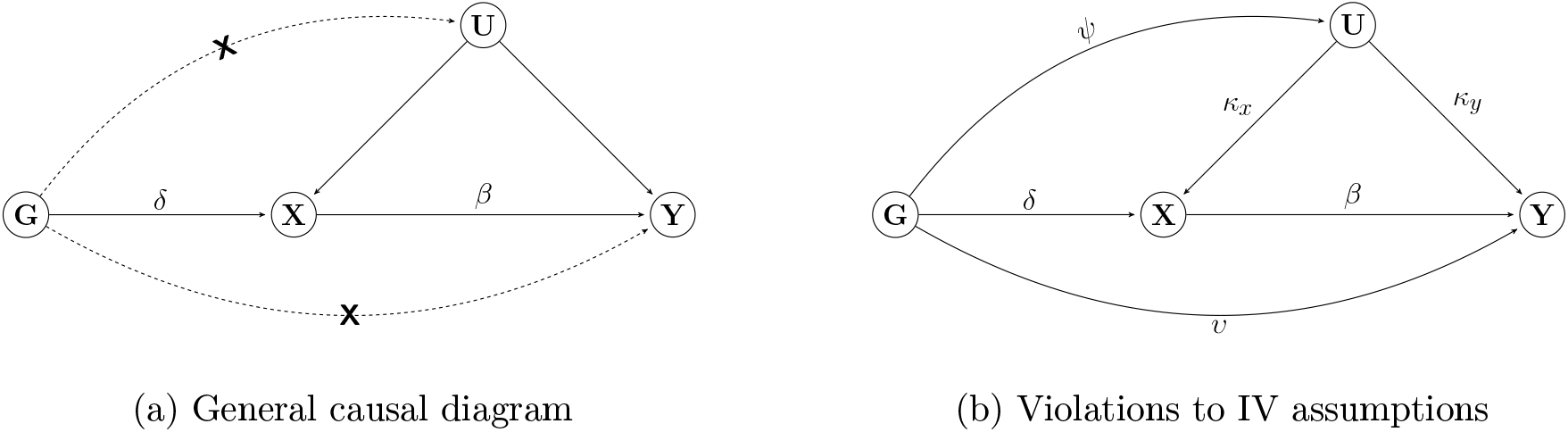
Causal diagrams representing the hypothesized relationship between genetic instrument (G), exposure (X), outcome (Y) and all unmeasured variables (U) which confound *X* and *Y*. *β* is the causal effect of X on Y. (a) *δ* is the genetic effect on X. Dashed lines and crosses indicate violations of the standard IV assumptions which can lead to bias. (b) Genetic instruments have a direct effect on Y (*v*), a phenomenon known as horizontal pleiotropy and a violation of IV3. Genetic instruments have a direct effect on U (*Ψ*), violation of IV2 and an example of horizontal pleiotropy that violates the InSIDE assumption.

Two-sample summary data MR is a design that derives causal effect estimates with summary statistics obtained from two separate samples - one supplying the Single Nucleotide Polymorphism (SNP)-exposure associations and the other supplying the SNP-outcome associations [3–6] - a SNP being the most common type of genetic variation in the genome. If the chosen SNPs are valid IVs, and the causal effect of a unit increase in *X* on the mean value or risk of *Y* is approximately linear in the local region of *X* predicted by these variants [7] then a simple inverse-variance weighted (IVW) meta-analysis of SNP-specific causal estimates provides an approximately unbiased estimate of this average causal effect. If sufficient heterogeneity exists between the MR estimates across a set of variants, this suggests evidence for violation of the IV assumptions. This could be due to assumption IV1 being only weakly satisfied by the genetic variants (i.e. weak instrument bias) [8, 9]. It is however more problematic when the heterogeneity is caused by violations of assumptions IV2 and IV3 [10, 7]. The latter violation is commonly known as "horizontal pleiotropy" [11], and hereafter referred to as pleiotropy for simplicity. Pleiotropy does not necessarily lead to biased causal effect estimates if it is balanced, in the sense that the average pleiotropic bias across SNPs is zero and the weight each SNP receives in the analysis is also independent of its pleiotropic effect. This latter condition is referred to as the Instrument Strength Independent of Direct Effect (InSIDE) assumption [12, 13]. However, this assumption is itself unverifiable.

Methods have been developed that are naturally robust to pleiotropy and InSIDE violation. For example, the weighted median estimator [14] provides a consistent estimate if 50% of the SNPs are valid IVs (or not pleiotropic). Similarly, mode-based estimation strategies focus on identifying the largest subset of variants yielding a homogeneous causal estimate, and are consistent when this set is made up of valid IVs [15, 16]. These approaches do not make any assumptions about the nature of the pleiotropy for invalid SNPs - they could violate InSIDE or not. Other approaches, such as MR-PRESSO [17] and Radial MR [8] attempt to detect and remove SNPs that are deemed responsible for bias and heterogeneity in an MR-analysis, however they assume the remaining SNPs satisfy InSIDE. Finally, the Robust Adjusted Profile Score (MR-RAPS) [9] uses an adjusted profile likelihood, which penalizes outlying (and hence likely pleiotropic) SNPs using a robust loss function.

In this paper we develop a method for pleiotropy robust MR analysis with two-sample summary data using the general framework of Bayesian Model Averaging (BMA) [18]. We adapt this general approach to the summary data setting where the SNPs are uncorrelated but potentially pleiotropic. Our approach uses the profile likelihood of MR-RAPS [9] as a basis for efficiently modelling the summary data in the presence of weak instrument bias and pleiotropy, but with the addition of an indicator function to denote whether an individual SNP is included or disregarded in the model. We develop a Metropolis-Hastings BMA algorithm to intelligently search the space models defined by all possible SNP subsets (i.e ≈ 2^*L*^ in the case of *L* SNPs) in order to decide which SNPs to include in the identified set of valid IVs within a given iteration of the markov chain. The derived posterior distribution is therefore averaged across all selected SNP combinations. We call our method BayEsian Set IDEntification Mendelian randomization (BESIDE-MR). BESIDE-MR aims to find the largest set of variants that furnish consistent, homogeneous estimates of causal effect, but accounts for model uncertainty, due to the selection of different instrument sets, which we will show is important for preserving the coverage of resulting MR estimates. Our one parameter BESIDE-MR model is robust to a small proportion of invalid SNPs, but is inadequate when a large proportion of SNPs are invalid. To address this case we extend MR-BESIDE to a two parameter model.

In Section 2 we introduce the methodology behind our one parameter model and in Section 3 assess its performance in Monte-Carlo simulations. In Section 4, we introduce and assess the performance of the two-parameter model extension. In Section 5, we apply both approaches to investigate the causal role of high density lipoprotein cholesterol (XL.HDL.C) on the risk of age related macular degeneration (AMD) using data from the 2019 MR Data Challenge [19]. We conclude with a discussion and point to further research.

## 2 Method

### 2.1 Description of the general model

Suppose that we have data from an MR study consisting of N individuals, where for each subject *k* we measure *L* independent genetic variants (*G*_*k*1_ ... *G*_*kL*_), an exposure (*X*_*k*_) and an outcome (*Y*_*k*_). *U*_*k*_ represents the shared residual error between *X* and *Y* due to confounding, which we wish to overcome using IV methods. To estimate the average causal effect, we assume the following linear structural models [20] for *U*, *X* and *Y* consistent with Figure 1b:

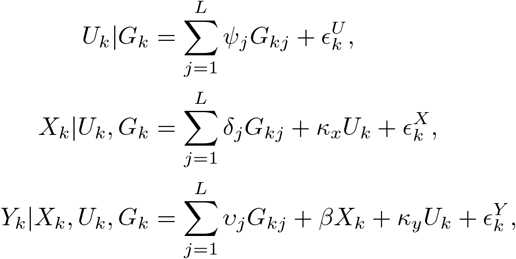

where 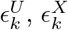 and 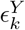 are mean zero independent error terms for *U*, *X* and *Y* respectively. See Appendix A.1 for summary of assumption required for the estimation of the average causal effect. From these structural models we can derive the approximate reduced form models for the *G-X* and *G-Y* associations for SNP *j* :

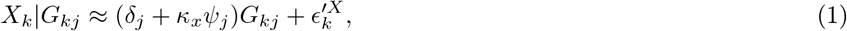

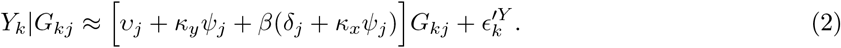

We use ‘approximate’ here because the error terms 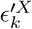 and 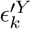 not strictly constant or mutually independent - the *j*th residual error term in fact contains common contributions from all other genetic variants not equal to *j*. This approximation is very accurate in most settings because the genetic variants combined make a very small contribution to the total residual error in each model (e.g. typically of the order of 1-2%) and the marginal coefficients are estimated from genome-wide association studies (GWAS) that usually have sample size of hundreds of thousands ([21]). Under this assumption the following models can then be justified for summary data estimates of the 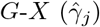 and 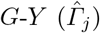 associations gleaned from fitting (1) and (2):

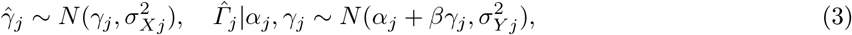

Here, *α*_*j*_ = *v*_*j*_ + *κ*_*y*_*Ψ*_*j*_, and *γ*_*j*_ = *δ*_*j*_ + *κ*_*x*_*Ψ*_*j*_. Under Model (16) it is assumed that the first study provides 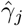 and standard errors *σ*_*Xj*_, and a second study, independent from the first, provides 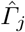 and standard errors *σ*_*Yj*_. Both the standard errors are assumed to be fixed and known. As the two studies are independent, we assume that the uncertainty in 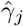 is independent of the uncertainty in 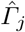. Model (16) also implies that SNPs are independent, independent SNPs can be found by performing linkage disequilibrium (LD) clumping in publicly available tools such as PLINK [22] and MR-BASE [23]. The two-sample design implicitly assumes that SNP *j* associations have identical associations in both studies as they are sampled from the same population. See Appendix A for formalised and further justification of the underlying assumptions made to estimate the average causal effect via two-sample approach.

The individual Wald ratio estimand for SNP *j* from Model (16) is then

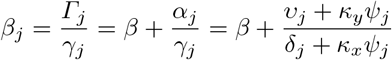

From this we see that to reduce the bias of *β*_*j*_ of SNP *j*, *γ*_*j*_, the instrument strength needs to be large. And/or *α*_*j*_, the amount of pleiotropic effect, either from SNP’s direct effect on *Y* (*v*_*j*_) or through *U* (*κ*_*y*_*Ψ*_*j*_), is close to zero. Under Model (16), invalid SNPs can be put into two classes:

**–** InSIDE respecting pleiotropy, *v*_*j*_ ≠ 0 but *Ψ*_*j*_ = 0
**–** InSIDE violating pleiotropy, *v*_*j*_ ≠ 0 and *Ψ*_*j*_ ≠ 0.

InSIDE violation occurs in the last case because instrument strength and pleiotropic effects are functionally related due to a shared *Ψ*_*j*_ component, so that the sample covariance 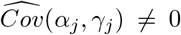. For the case of InSIDE respecting pleiotropy we are able to assume the sample covariance is approximately zero for a sufficient number of instruments, since *v*_*j*_ and *δ*_*j*_ are imagined to be themselves generated via independent processes [7]. In Appendix B, we show, under the simplifying assumption that the SNP-outcome standard errors are approximately constantand *κ*_*x*_ = *κ*_*y*_ = 1, when 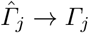 and 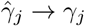 as N → ∞, the approximate bias term for IVW estimator is,

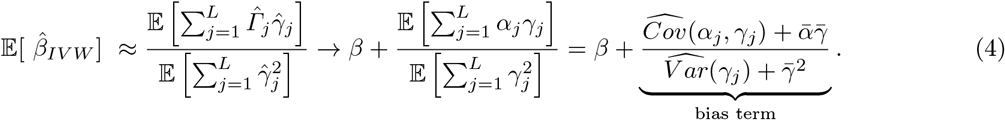

If all SNPs are pleiotropic, but have mean zero (ᾱ=0) and satisfy the InSIDE assumption 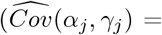 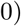, then the standard IVW provides an unbiased estimate of *β*. MR-Egger regression is an extension of IVW that can work under the InSIDE assumption even if ᾱ = 0, which is referred to as ‘directional’ pleiotropy. It does this by estimating an intercept parameter in addition to the causal slope parameter. However, its estimates are generally very imprecise and it is not invariant to allele recoding [24]. Lastly, it can not separate directional pleiotropy satisfying InSIDE from balanced pleiotropy violating InSIDE, as the intercept reflects the numerator of the bias term, which is a combination of both. This motivates the use of methods that can attempt to detect and down-weight a small number of variants that may be responsible for either InSIDE violation or directional pleiotropy so that, for the remainder of SNPs left, Model (16) holds with only InSIDE respecting balanced pleiotropy. This approach we will initially pursue for BESIDE-MR, which also have been taken by others [9, 17]. Another further advantage by not estimating an intercept term, BESIDE-MR will be invariant to allele recoding, as opposed to MR-Egger.

### 2.2 Bayesian Model Averaging over the summary data model

We are interested in searching over the space of all possible models defined by each of the 2^*L*^ subsets in the entire summary data. Let *I* = (*I*_1_, ..., *I*_*L*_) be the *L*-length indicator vector denoting whether SNP *G*_*j*_ is included (*I*_*j*_ = 1) or not (*I*_*j*_ = 0) in the model. We want to ‘force’ our data to conform to Model (16) with the additional assumption that *α*_*j*_ ~ *N*(0, *τ*^2^). The parameters of interest are then *θ* = (*β, τ*^2^, *I*) and with data, *D*, that consists of 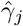 and 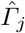, with their standard errors *σ*_*Xj*_ and *σ*_*Yj*_ respectively. Then the joint posterior is

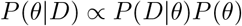

where *P*(*D|θ*) is the likelihood and *P*(*θ*) is user specified prior for each of the parameters. We use a random walk Metropolis-Hastings (M-H) algorithm for updating the model parameter values, for the specific details see Appendix C. For a given iteration of the markov chain, the selection of instruments is conditional on the likelihood of the data and the given priors. After the markov chain has been sufficiently explored, we can obtain posterior distributions for the model parameters and the posterior probability that each individual SNP is valid. This method has been found in individual-level data to reduce bias from many weak instruments [25, 26] and highly correlated instruments [27]. It has been previously shown that using a small number of SNPs for two-sample MR can lead to large violations of the InSIDE assumption by chance (see Figure A.1 in [7]). Small SNP numbers also make estimation of the pleiotropy variance very imprecise. Therefore, we have restricted the M-H algorithm to explore models that have at least 5 instruments. Given that the BESIDE-MR model is weak-instrument robust, it will almost always be possible to include a sufficient number of instruments because it is not necessary to select only ‘genome-wide significant’ SNPs - a weaker selection threshold can be used.

**The profile score likelihood** For *P*(*D*|*θ*), we use the profile log-likelihood score derived by [9].

Specifically this is the likelihood for (*β, τ*^2^) given the data 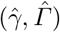 profiled over the parameters *γ*_1_, ..., *γ*_*L*_. After the incorporation of our indicator vector *I*, the log-likelihood is modified to

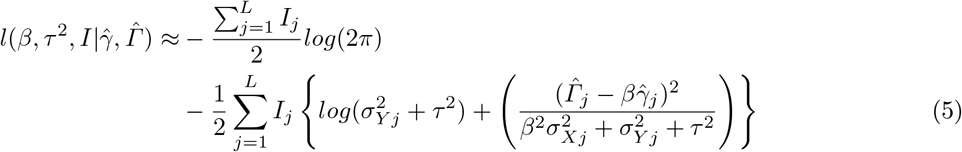

As shown by the derivation in Appendix D, this likelihood allows for heterogeneity due to pleiotropy via *τ*^2^, and weak instruments, via 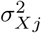. If we consider that the existing set of instruments have a small *τ*^2^, then the likelihood will increase if introducing a new instrument does not lead to a sufficiently large increase the pleiotropy variance, in which case it may decrease. Hence, our BMA algorithm will naturally give more weight to *I*-vectors that include large sets of instruments with homogeneous causal effect estimates. This property is reminiscent of the ZEro Modal Pleiotropy Assumption (ZEMPA) [15] or the plurality rule that defines the two-stage hard thresholding (TSHT) approach of Guo *et al.* [28]. However, the TSHT approach explicitly aims to isolate the largest set of ‘valid’ instruments and base all inference on this single set, which is equivalent to giving a single *I*-vector a weight of 1 and all other vectors a weight of zero. BESIDE-MR is less aggressive, allowing as many distinct *I*-vectors as are supported by the data to be given weight in the analysis. This feature properly accounts for model uncertainty. Indeed, as subsequent simulations will demonstrate, this yields causal estimates and standard errors that are less prone to under-coverage than methods which incorporate instrument selection or penalization.

One such method of penalization, also proposed by Zhao *et al.* [9], is MR-RAPS. Instead of being based on likelihood function (5) which uses standard least squares (or *L*_2_ loss) plus the addition of our indicator function, it uses a robust L1 function such as Huber or Tukey loss. This enable the contribution of large outliers to be penalized (i.e. reduced) compared to *L*_2_ loss. Our use of the standard profile likelihood can be viewed as an alternative way to achieving the robustness of MR-RAPS, by averaging over multiple instruments sets and where more weight is given to homogeneous SNP sets. For this very reason, convergence is an essential part of BESIDE-MR implementation to ensure that all plausible models have been explored. The profile likelihood is particularly well suited to a Bayesian implementation because it enables heterogeneity due to weak instrument bias and pleiotropy to be accounted for, whilst only having to update three parameters (*β*, *τ* and *I*). Generally, a standard Bayesian formulations requires an additional *L* parameters (*γ*_1_, ..., *γ*_*L*_) to be updated, (e.g. see Thompson *et al.* [29]).

BMA implementations tends to favour parsimonious models, i.e. models with fewer variables [18], therefore, to explore the sensitivity of our BMA procedure to the average number of SNPs included in the model, we include a penalization term within likelihood function (5);

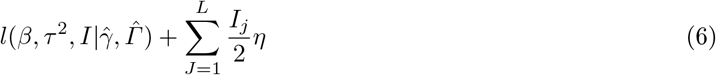

*η* will dictate which size models BMA should explored the most; setting a large positive *η*, the likelihood will increase with number of instruments, then BMA will favour models with many instruments. And hence for negative *η*, BMA will favour models with fewer instruments. We will assume *η* to be zero throughout the simulations, but explore ranges of *η* as sensitivity analysis for the real data example in Section 5.1.

### 2.3 Choice of priors

In general we encourage the construction of priors to be based on previous epidemiological study or biological knowledge. For the purpose of elucidating our approach, we will use priors that ensure efficient mixing and rapid convergence. For the causal effect parameter *β*, we use a zero centered normal prior *P*(*β*). For the pleiotropy variance (*τ*^2^) we use a gamma prior *P*(*Prec*) for the precision, where *Prec* = 1*/τ*^2^. For the indicator function prior, we will assume an uninformative Bernoulli prior *P*(*I*) with probability 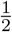 for all *I*_*j*_.

### 2.4 An alternative implementation

It is well known that the estimation of *τ*^2^ is challenging, even within a classical framework, as its maximum likelihood estimate is not consistent, see Section 4 of Zhao *et al.* [9] for further discussion. Therefore, we propose an alternative implementation of our M-H algorithm in which a plug-in estimate for *τ*^2^ is substituted at each iteration. For simplicity, we chose to use the closed-form DerSimonian-Laird estimate for *τ*^2^ [30]. In Appendix C, we describe how the M-H algorithm is modified to implement this alternative approach. Hereafter, we will refer to the first method as the ‘full Bayesian’ approach and this latter method as the DerSimonian-Laird (DL) approach.

## 3 Monte Carlo simulation

### 3.1 Simulation strategy

We simulate two-sample summary MR data sets with *L*=50 instruments from Model (16). Motivated by recent genetic studies [31, 32], four scenarios are considered;

1. all instruments are strong and invalid instruments have balanced pleiotropy,
2. all instruments are weak and invalid instruments have balanced pleiotropy,
3. all instruments are strong and invalid instruments have directional pleiotropy,
4. all instruments are weak and invalid instruments have directional pleiotropy.

The strength of the instruments is determined by mean F-statistic 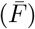 over all instruments. The pleiotropic effect of invalid instruments, *α*_*j*_, is simulated from a normal *N*(*μ*_*α*_, *σ*_*α*_) distribution, where zero and non-zero *μ*_*α*_ gives balanced and directional pleiotropy respectively, as shown in Table 1. Note that, whilst scenarios 3 and 4 are referred to as directional pleiotropy, this could equally be thought of as InSIDE violating pleiotropy, as illustrated in Equation (4). Within each scenario, 0% to 100% (at 20% intervals) of the L SNPs are simulated as invalid instruments. We first compare our approach with the standard IVW method, MR-APS and MR-RAPS. The latter two are the classical counterparts that our approach sits between. Specifically, MR-APS is the MR-RAPS with a standard L2 loss function as opposed to Huber or Tukey loss. We monitor the mean bias of the causal parameter estimate and the coverage (for BESIDE-MR the bias is taken with respect to the mean of the posterior distribution of *β* and the coverage is calculated from its credible interval). For BESIDE-MR only, we also give the average difference in the posterior probability of inclusion, *Δ*(*PPI*), to show how often each SNP can be correctly assigned to its true set. We also report the weak instrument bias corrected exact *Q*-statistic [8] to measure the amount of heterogeneity in our simulated data. See details of the simulation strategy in Appendix E.

**Table 1:** Summary of simulation scenarios.

| Scenario | Type of pleiotropy | $\bar{F}$ | pleiotropic effect ($\alpha_j$)<br>of invalid instruments |
| --- | --- | --- | --- |
| 1 | Balanced | 100 | $N(0, 0.04)$ |
| 2 | Balanced | 10 | $N(0, 0.04)$ |
| 3 | Directional | 100 | $N(0.05, 0.04)$ |
| 4 | Directional | 10 | $N(0.05, 0.04)$ |

From the convergence test, our algorithm functions effectively with 50,000 iterations with 10,000 burn-ins. For each simulated dataset of 50 instruments, DL and fully Bayesian implementation took 5 and 7 seconds to converge respectively on a standard desktop computer, however number of iterations needs to increase with more instruments to ensure convergence, as it takes longer for BESIDE-MR to explore all feasible models out of a potential 2^*L*^. In rare occasions, we removed results simulations where the BESIDE-MR model had failed to converge after the set number of iterations (see results from the convergence test in Appendix E.2).

### 3.2 Results

Table 2 shows the results. Under Scenario 1, all methods deliver approximately unbiased estimates. The IVW, MR-APS and MR-RAPS estimators achieve nominal coverage when there are no pleiotropic instruments. However, as the proportion of pleiotropic instruments (and hence the heterogeneity) increases, their coverages can drop substantially, with the MR-APS and MR-RAPS estimators most affected. BESIDE-MR has conservative coverage under no heterogeneity (due to the absence of invalid instruments) but maintains far better coverage when this increases. The general pattern remains the same for weaker instruments (Scenario 2), even with many more weak instruments (*L*=100, results shown in Appendix E.4). In Scenarios 3, all the approaches deteriorate with increasing number of invalid instruments, but BMA has consistently the least bias and best coverage throughout. In Scenario 4, the IVW estimator is seemingly least biased, due to weak instrument bias cancelling out some of the pleiotropic bias. With 40% and 60% invalid instruments, full Bayesian BESIDE-MR struggled to converge within 50,000 iterations in a small number of cases.

**Table 2:** Evaluation criteria for different types of pleiotropy and instrument strength (Table 1. 50 instruments in total. True *β* is 0.05. No. inv., Number of invalid instrument(s); *Q*, Q-statistics with exact weights; DL est., DL estimate; Full Bayes., full Bayesian; Bias, mean bias; Cover., coverage.).

| No. inv. | $Q$ | IVW | | DL est. | | Full Bayes. | | MR-APS | | MR-RAPS | |
| --- | --- | --- | --- | --- | --- | --- | --- | --- | --- | --- | --- |
|  |  | Bias | Cover. | Bias | Cover. | Bias | Cover. | Bias | Cover. | Bias | Cover. |
| Scenario 1 |  |  |  |  |  |  |  |  |  |  |  |
| 0 | 49.0 | -0.001 | 96.40 | -0.000 | 97.50 | 0.000 | 98.10 | -0.000 | 94.40 | -0.000 | 94.00 |
| 10 | 57.9 | -0.001 | 93.20 | 0.000 | 97.50 | 0.000 | 97.70 | -0.000 | 89.50 | -0.000 | 92.10 |
| 20 | 66.4 | -0.001 | 90.80 | -0.000 | 95.40 | -0.000 | 94.60 | -0.000 | 83.90 | -0.000 | 87.30 |
| 30 | 75.5 | -0.000 | 88.30 | 0.001 | 94.20 | 0.001 | 92.00 | 0.001 | 77.30 | 0.001 | 80.80 |
| 40 | 84.0 | -0.001 | 86.80 | -0.000 | 95.80 | -0.000 | 90.70 | 0.001 | 76.60 | 0.001 | 77.60 |
| 50 | 91.9 | 0.000 | 85.40 | 0.000 | 94.80 | 0.001 | 86.60 | 0.002 | 70.40 | 0.001 | 72.90 |
| Scenario 2 |  |  |  |  |  |  |  |  |  |  |  |
| 0 | 48.7 | -0.018 | 33.40 | -0.001 | 97.10 | 0.002 | 96.10 | -0.000 | 93.90 | -0.000 | 92.90 |
| 10 | 54.4 | -0.019 | 37.50 | -0.000 | 97.10 | 0.005 | 93.70 | 0.003 | 91.80 | 0.003 | 92.10 |
| 20 | 59.2 | -0.018 | 41.70 | 0.001 | 96.70 | 0.008 | 90.50 | 0.006 | 88.00 | 0.006 | 89.10 |
| 30 | 64.0 | -0.018 | 44.60 | 0.001 | 96.70 | 0.011 | 87.80 | 0.009 | 83.20 | 0.008 | 84.90 |
| 40 | 68.8 | -0.018 | 46.50 | 0.001 | 95.60 | 0.014 | 80.20 | 0.012 | 72.50 | 0.011 | 75.70 |
| 50 | 73.9 | -0.019 | 47.80 | 0.002 | 94.60 | 0.017 | 73.40 | 0.015 | 68.80 | 0.015 | 70.10 |
| Scenario 3 |  |  |  |  |  |  |  |  |  |  |  |
| 0 | 49.0 | -0.001 | 96.40 | -0.000 | 97.50 | 0.000 | 98.10 | -0.000 | 94.40 | -0.000 | 94.00 |
| 10 | 69.0 | 0.011 | 75.60 | 0.007 | 92.80 | 0.007 | 92.70 | 0.013 | 61.30 | 0.009 | 75.80 |
| 20 | 84.1 | 0.024 | 35.20 | 0.018 | 71.90 | 0.016 | 70.00 | 0.027 | 20.20 | 0.021 | 33.60 |
| 30 | 92.0 | 0.037 | 11.80 | 0.032 | 38.20 | 0.031 | 36.10 | 0.039 | 4.70 | 0.035 | 7.90 |
| 40 | 96.1 | 0.051 | 1.40 | 0.049 | 9.30 | 0.049 | 9.70 | 0.054 | 0.10 | 0.052 | 0.40 |
| 50 | 95.2 | 0.064 | 0.30 | 0.066 | 1.50 | 0.067 | 1.50 | 0.068 | 0.00 | 0.067 | 0.00 |
| Scenario 4 |  |  |  |  |  |  |  |  |  |  |  |
| 0 | 48.7 | -0.018 | 33.40 | -0.001 | 97.10 | 0.002 | 96.10 | -0.000 | 93.90 | -0.000 | 92.90 |
| 10 | 58.8 | -0.011 | 69.77 | 0.007 | 95.60 | 0.015 | 79.00 | 0.018 | 66.30 | 0.016 | 71.70 |
| 20 | 64.5 | -0.003 | 84.70 | 0.017 | 84.60 | 0.028 | 46.20 | 0.035 | 23.70 | 0.034 | 29.60 |
| 30 | 66.5 | 0.006 | 82.60 | 0.028 | 64.60 | 0.040 | 21.70 | 0.050 | 5.10 | 0.048 | 7.00 |
| 40 | 66.2 | 0.014 | 70.10 | 0.040 | 35.60 | 0.049 | 9.90 | 0.064 | 0.40 | 0.063 | 0.60 |
| 50 | 65.3 | 0.022 | 53.90 | 0.050 | 18.90 | 0.057 | 5.20 | 0.075 | 0.10 | 0.074 | 0.10 |

*Δ*(*PPI*) in Figure 2 demonstrates BESIDE-MR’s ability to distinguish valid instruments in the presence of invalid instruments for Scenarios 1 and 3. For valid SNPs to be correctly identified we want *Δ*(*PPI*) to be large and positive. This difference should of course be zero when there are no invalid instruments. Under Scenario 1 this difference is maximised (i.e. we get the best discrimination) when there are 20% invalid instruments, this difference steadily decreases to half its value as the number of invalid instruments increases further. Under Scenario 3 we see a smaller and more constant difference across different proportions of invalid instruments, indicating that BESIDE-MR generally struggles to deal with directional/InSIDE violating pleiotropy across a substantial proportion of invalid SNPs. There is still a difference in *PPI* between valid and invalid instruments, however the discrimination is worse for weak instruments.

**Fig. 2:**
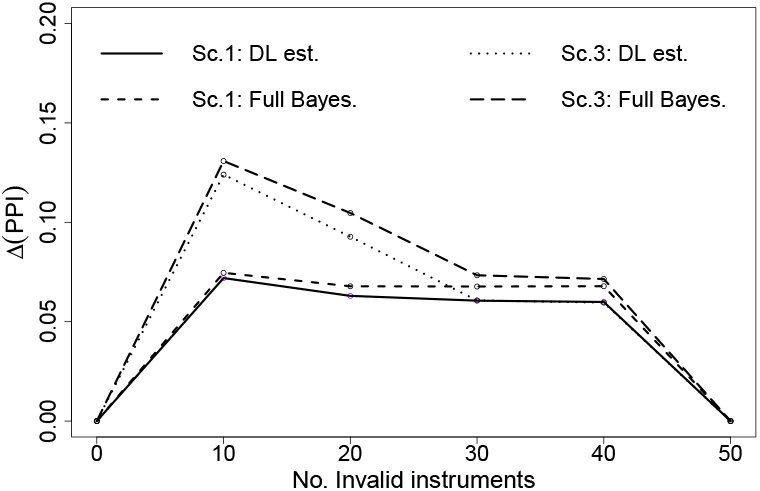
The difference in posterior probabilities of inclusion (*Δ*(*PPI*)) between valid and invalid instruments for balanced and directional pleiotropy (Scenario 1 and 3 respectively). On the x-axis is the number of invalid/pleiotropic instruments, and the y-axis is the average difference in PPI in valid and invalid instruments set, *Δ*(*PPI*), over 1,000 simulations. As shown by legend within plot, the lines denotes results from different implementation of BESIDE-MR within each scenario.

Additional simulations were performed to investigate the effect of different patterns of heterogeneity, on *Δ*(*PPI*). We find that the discrimination is best with small numbers of highly pleiotropic SNPs, and the worst with large numbers of weakly pleiotropic SNPs. However, the algorithm maintains its reliability even in this case. For further details see Appendix E.5 for the results.

## 4 An extended two-parameter BMA model for InSIDE violation

The one (causal) parameter BESIDE-MR model introduced thus far assumed that most SNPs were valid under the InSIDE assumption, but a small proportion could be invalid under InSIDE. We now consider the use of an extended model to account for the more extreme case where a large proportion of SNPs may be pleiotropic, and in violation of InSIDE (Figure 1b). In this case, the standard one parameter BESIDE-MR model cannot easily identify and remove the invalid SNPs, they must instead be formally modelled with an additional slope parameter. To motivate this approach we use the same underlying data generating Model (16). Suppose that we have two different groups of invalid instruments: in the first group, *S*_1_, the SNPs exhibit balanced pleiotropy under the InSIDE assumption, but still collectively identify the true causal effect, *β*. For illustrative purposes, suppose now that all of the remaining instruments are in a set *S*_2_, where the InSIDE assumption is perfectly violated (that is, the correlation between the SNP-exposure association and the pleiotropic effect is 1). Using the bias formulae in Equation (4), the set of SNPs in *S*_2_ identify a distinct, biased version of the causal effect (*β* + 1). This data generating model would give rise to two clusters or slopes in the data, which motivates our extended two-parameter version of BESIDE-MR.

### 4.1 A modified BMA algorithm

Under the data generating Model (16), we further assume that the pleiotropic effects for valid SNPs in *S*_1_ are generated from a 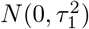 distribution and the invalid SNPs in *S*_2_ are from a 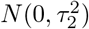 distribution. Allowing these SNPs to violate InSIDE, and therefore identify a different slope parameter, our total parameter space is modified to 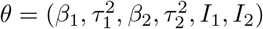, with likelihood:

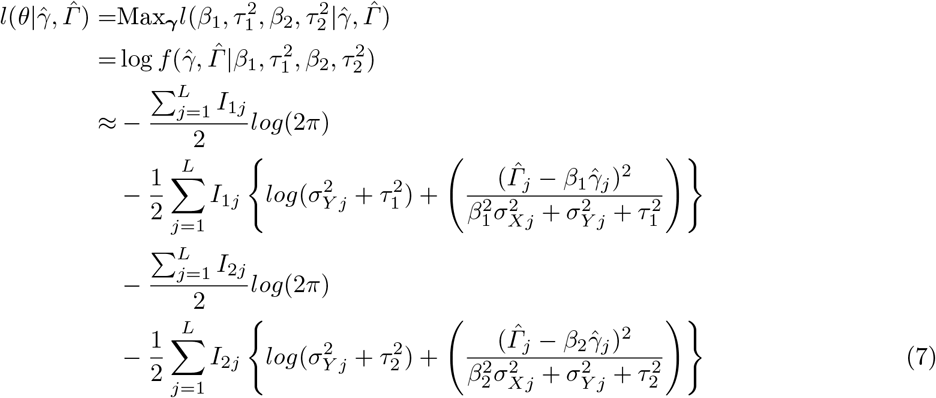

where the indicator functions *I*_1*j*_ and *I*_2*j*_ denote whether a SNP *j* is included in *S*_1_ or *S*_2_. We impose the condition that *I*_1*j*_ + *I*_2*j*_ ≤ 1, which means that, at a given iteration of our BMA algorithm a SNP is either in *S*_1_ (*I*_1*j*_ = 1, *I*_2*j*_ = 0), *S*_2_ (*I*_1*j*_ = 0, *I*_2*j*_ = 1) or in neither *S*_1_ or *S*_2_ (*I*_1*j*_ = *I*_2*j*_ = 0), which we give the label *S*_0_. This gives the model the flexibility to assign a SNP to either *S*_1_ or *S*_2_, or remove it from the model completely by assigning it to *S*_0_. In Appendix F, we give further details on the M-H algorithm to update the parameter space of this extended model.

The log-likelihood with the addition of two model complexity penalisation terms is then;

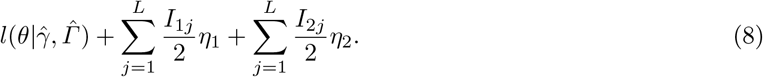

Same as in Section 2.2, we set *η*_1_ = *η*_2_ = 0 for the simulations, but vary the values as sensitivity in the applied example.

### 4.2 Simulation study

Two-sample summary data are simulated with 50 SNPs under balanced pleiotropy but with progressively larger proportion of SNPs maximally violating the InSIDE assumption. This changes the proportion of SNPs that are in set *S*_1_ and *S*_2_. These data are simulated under a strong instrument scenario (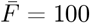, Scenario 5) and a weaker instrument scenario (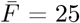, Scenario 6). For precise details of the simulation parameters see Table 3. We also explore the performance of our two-parameter model under balanced pleiotropy with weak and strong instruments when there is no InSIDE violation. That is, under Scenario’s 1 and 2. This means that all SNPs are effectively in set *S*_1_ and the data can be explained with a single causal slope parameter, not two. The full results are shown in Table 4 where we report the bias, coverage and mean Q-statistic with exact weights of all approaches across 1,000 simulations, as before. For BESIDE-MR, 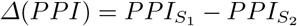 is also reported for SNPs truly in *S*_1_ and *S*_2_.

**Table 3:** Summary of InSIDE simulation scenarios

| Scenario | $\bar{F}$ of $S_1 : S_2$ | Type of pleiotropy | $S_1$ | $S_2$ |
| --- | --- | --- | --- | --- |
| 5 | 100:100 | Balanced | $\psi_j = 0,$<br>$v_j \sim N(0, 0.04),$<br>$\delta_j \sim U(0.34, 1.1),$<br>$\sigma_{X_j} \sim U(0.06, 0.095),$<br>$\beta_1 = \beta$ | $\psi_j \sim U(0.34, 1.1),$<br>$v_j = 0,$<br>$\delta_j = 0,$<br>$\sigma_{X_j} \sim U(0.06, 0.095),$<br>$\beta_2 = \beta + 1$ |
| 6 | 25:25 | Balanced | $\psi_j = 0,$<br>$v_j \sim N(0, 0.04),$<br>$\delta_j \sim U(0.34, 1.1),$<br>$\sigma_{X_j} \sim U(0.06, 0.4),$<br>$\beta_1 = \beta$ | $\psi_j \sim U(0.34, 1.1),$<br>$v_j = 0,$<br>$\delta_j = 0,$<br>$\sigma_{X_j} \sim U(0.06, 0.4),$<br>$\beta_2 = \beta + 1$ |

**Table 4:** Evaluation criteria for estimating two causal parameter. 50 instruments in total. The true *β* is 0.05. *S*_1_ and *S*_2_ are InSIDE respecting and violating set respectively. Est., estimator; Inst., instrument(s); *Q*, exact Q-statistics; DL est., DL estimate; Full Bayes., Full Bayesian.

| Est. | Inst. $S_1 : S_2$ | Q | | mean bias | | median bias | | coverage | |
| --- | --- | --- | --- | --- | --- | --- | --- | --- | --- |
| | | $S_1$ | $S_2$ | $\beta_1$ | $\beta_2$ | $\beta_1$ | $\beta_2$ | $\beta_1$ | $\beta_2$ |
| **Scenario 1** ($\beta_1 = \beta_2 = \beta$) | | | | | | | | | |
| DL est. | 50:0 | 60.2 | - | 0.001 | 0.001 | 0.001 | 0.001 | 100.0 | 99.8 |
| Full Bayes. | 50:0 | 60.2 | - | 0.001 | 0.001 | 0.001 | 0.001 | 99.7 | 99.5 |
| **Scenario 5** ($\beta_1 = \beta, \beta_2 = \beta + 1$) | | | | | | | | | |
| DL est. | 40:10 | 73.5 | 10.9 | 0.007 | -0.876 | 0.001 | -0.995 | 99.4 | 14.6 |
|  | 30:20 | 55.1 | 23.8 | 0.003 | -0.079 | 0.001 | -0.013 | 95.9 | 92.3 |
|  | 25:25 | 43.9 | 30.3 | 0.005 | -0.009 | 0.001 | -0.008 | 93.9 | 96.7 |
|  | 20:30 | 35.3 | 36.9 | 0.053 | -0.006 | 0.004 | -0.006 | 91.0 | 95.6 |
|  | 10:40 | 16.5 | 49.1 | 0.906 | -0.008 | 0.988 | -0.005 | 11.1 | 85.5 |
| Full Bayes. | 40:10 | 73.5 | 10.9 | 0.076 | -0.287 | 0.003 | -0.027 | 84.0 | 69.0 |
|  | 30:20 | 55.1 | 23.8 | 0.230 | -0.218 | 0.008 | -0.009 | 69.4 | 76.2 |
|  | 25:25 | 43.9 | 30.3 | 0.248 | -0.182 | 0.011 | -0.008 | 67.9 | 79.7 |
|  | 20:30 | 35.3 | 36.9 | 0.254 | -0.122 | 0.013 | -0.002 | 66.8 | 86.1 |
|  | 10:40 | 16.5 | 49.1 | 0.225 | -0.041 | 0.017 | 0.003 | 62.4 | 95.4 |
| **Scenario 2** ($\beta_1 = \beta_2 = \beta$) | | | | | | | | | |
| DL est. | 50:0 | 58.3 | - | 0.002 | 0.002 | 0.002 | 0.002 | 100.0 | 100.0 |
| Full Bayes. | 50:0 | 58.3 | - | 0.004 | 0.004 | 0.004 | 0.003 | 99.9 | 99.9 |
| **Scenario 6** ($\beta_1 = \beta, \beta_2 = \beta + 1$) | | | | | | | | | |
| DL est. | 40:10 | 67.6 | 30.2 | 0.003 | -0.985 | 0.002 | -0.997 | 99.0 | 1.4 |
|  | 30:20 | 50.3 | 65.7 | 0.035 | -0.474 | 0.009 | -0.391 | 97.5 | 60.1 |
|  | 25:25 | 41.3 | 85.0 | 0.012 | -0.099 | 0.006 | -0.060 | 94.1 | 93.3 |
|  | 20:30 | 32.8 | 102.6 | 0.007 | -0.037 | 0.005 | -0.033 | 94.6 | 96.8 |
|  | 10:40 | 14.8 | 140.6 | 0.651 | -0.072 | 0.766 | -0.062 | 41.4 | 93.8 |
| Full Bayes. | 40:10 | 67.6 | 30.2 | 0.001 | -0.337 | 0.003 | -0.104 | 89.8 | 63.2 |
|  | 30:20 | 50.3 | 65.7 | 0.022 | -0.179 | 0.008 | 0.016 | 84.7 | 78.5 |
|  | 25:25 | 41.3 | 85.0 | 0.036 | -0.233 | 0.011 | 0.013 | 72.8 | 80.9 |
|  | 20:30 | 32.8 | 102.6 | 0.002 | -0.332 | 0.011 | 0.016 | 64.3 | 77.5 |
|  | 10:40 | 14.8 | 140.6 | 0.364 | -1.349 | 0.987 | -0.379 | 22.3 | 52.8 |

### 4.3 Results

For data generated under Scenario 1 and 2, and so in the complete absence of InSIDE-violating SNPs in set *S*_2_, our two slope model correctly identifies *β* and does not try to estimate a second effect, i.e. *β*_1_ = *β*_2_. When the data are generated under Scenario 5 we see that, when *S*_1_ and *S*_2_ have a similar number of instruments, both *β*_1_ and *β*_2_ can be estimated by the DL implementation of our two-parameter model. If the proportion of SNPs in either set is too small, then our algorithm tends to remove them completely and focus on estimating just one slope. The full Bayesian implementation returns mean posterior estimates that are median unbiased but not mean unbiased. This demonstrates a lack of convergence for some of simulated data, and indicates that longer iterations and a more sophisticated procedure for deciding on the tuning parameter may be required to properly fit the model.

When the data are generated with weaker instruments (Scenario 6), we see a degrading in the performance of all approaches. In particular, see that the effect is worst for *β*_2_. This is because, in our specific simulation, *β*_2_ is larger in magnitude than *β*_1_, which increases both the heterogeneity as measured by the exact *Q* statistic (see Equation 9 in Appendix E.4) and the absolute magnitude of weak instrument bias relative to that experienced when estimating *β*_1_. This adversely affected the coverage of the estimates. This heterogeneity is further exaggerated with weaker instruments 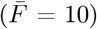, leading to our approach not being able to correctly assign instruments to either *S*_1_ or *S*_2_ (Appendix G.1). If this case is encountered in practice, we recommend use of the single slope model instead.

When applying the full Bayesian implementation of BESIDE-MR in Scenario 6, we noticed an important feature most prominent when there was a large imbalance in the relative sizes of *S*_1_ and *S*_2_. In this case, the M-H algorithm can switch from estimating the posterior for *β*_1_ to estimating the posterior for *β*_2_. This problem is referred to as “label switching” [33]. In our applied analysis in Section 5 we discuss this issue in more detail, and our proposal for addressing it.

Figure 3a gives further insight into how well the DL and full Bayesian implementations can correctly partition the SNPs into clusters. The x-axis shows the true ratio of SNPs in *S*_1_ and *S*_2_ and the y-axis shows *Δ*(*PPI*). For example, with instrument ratio 1:1, *Δ*(*PPI*) is positive for SNPs truly in *S*_1_ and negative for SNPs truly in *S*_2_. The DL estimate show that at ratio of 4:1 *Δ*(*PPI*) is similar for SNPs truly in *S*_1_ or *S*_2_, this is because as explained above, the DL approach more aggressively prefers to estimate 1 parameter only, and treating minority SNPs as outliers (e.g. assign to *S*_0_). By contrast, *Δ*(*PPI*) for the full Bayesian approach is much more constant across all ratios and are also consistently lower. When the *S*_1_:*S*_2_ ratio is balanced, both implementations correctly identified *S*_1_ and *S*_2_ instruments. However, due to “label switching”, both implementations struggles to identify *S*_1_ and *S*_2_ SNPs with weak instruments and larger proportion of *S*_2_ SNPs.

**Fig. 3:**
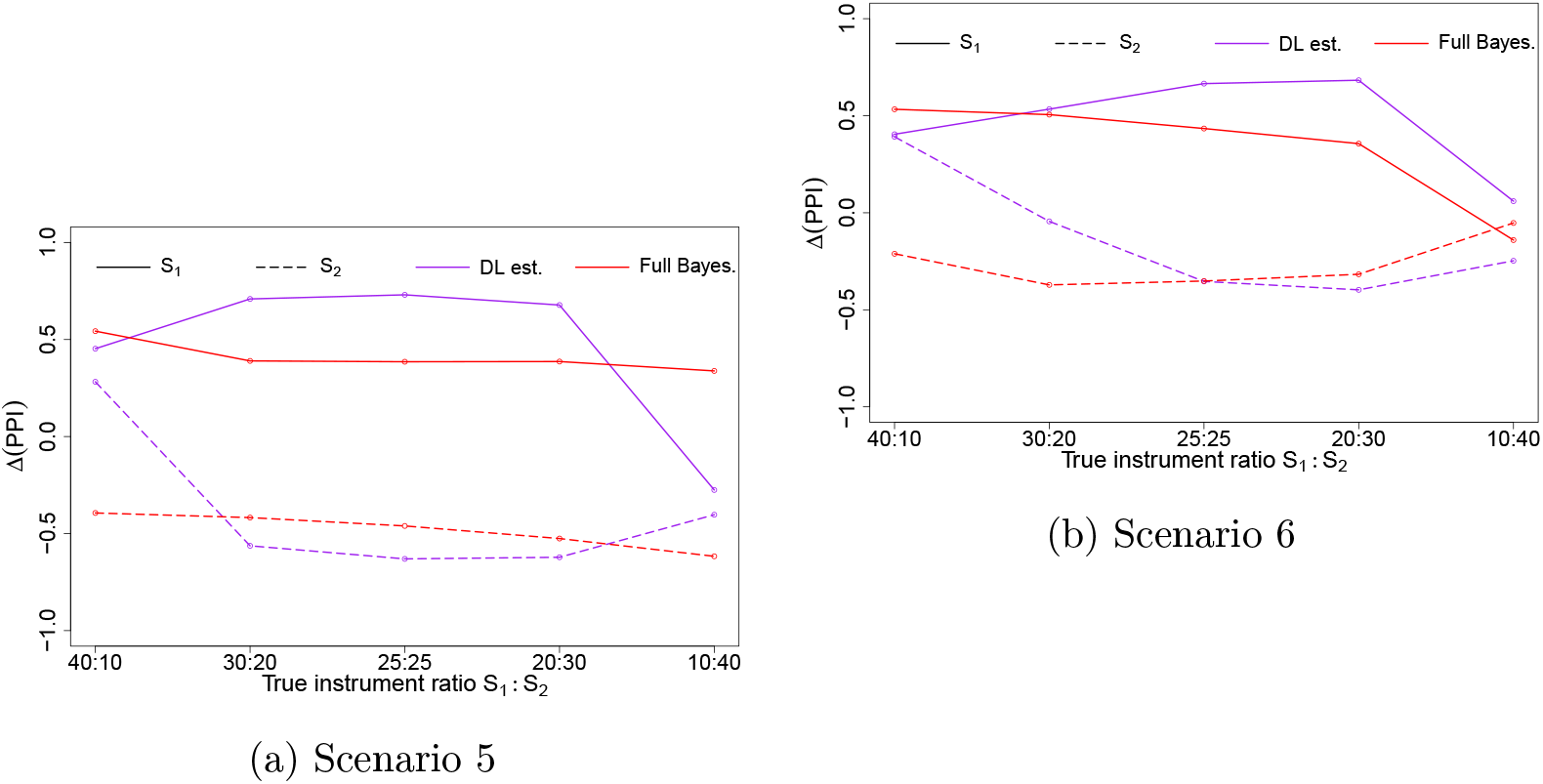
Identifying *S*_1_ and *S*_2_ instruments; their difference in inclusion probability (*Δ*(*PPI*)) for Scenario 5 (a) and Scenario 6 (b). The x-axis is the true ratio for number of instruments in each cluster (*S*_1_:*S*_2_), and the y-axis is *Δ*(*PPI*), averaged over 1,000 simulations. As shown by the legend within plot, the horizontal and dashed lines denotes *S*_1_ and *S*_2_ are BESIDE-MR classed *S*_1_ and *S*_2_ instruments respectively. The the purple and red colour denotes results from the 2 estimation approaches of BESIDE-MR.

If a SNP increases the overall heterogeneity (*τ*^2^) in either cluster, BESIDE-MR increasingly classes it as belonging to *S*_0_ (neither *S*_1_ or *S*_2_). Using a simulated example, Appendix Figure A.7, demonstrates that the further the SNP is from either of the slope line, the higher (darker in colour) the probability it belongs to neither clusters.

## 5 Applied example: Age-related macular degeneration and cholesterol

Age-related macular degeneration (AMD) is a painless eye-disease that eventually leads to vision loss. Recent GWAS have identified several rare and common variants located in gene regions that are associated with lipid levels [34], fuelling speculation as to whether the relationship is causal [35, 36]. To this end, a multivariable MR analysis was performed by Burgess *et al.* [37], which provided evidence to support a causal relationship between AMD and HDL cholesterol but not with LDL cholesterol and triglycerides. In follow up work, Zuber *et al.* [38] fitted a multivariable MR model using Bayesian model averaging, with a total of 30 separate lipid fraction metabolites acting as the intermediate exposures. Out of the 30, large particle HDL cholesterol (XL.HDL.C) had the highest inclusion probability as a risk factor for AMD.

Although multivariable MR approaches can remove bias due to pleiotropy via known pleiotropic pathways (in this case, other lipid fractions), they can be much more challenging to fit, especially when the correlation between the included exposures is high. For this reason we now revisit this data and use our univariate MR approaches to probe the causal relationship between XL.HDL.C and AMD.

To avoid selection bias, we selected 27 genetic variants as instruments from a separate sample, the METSIM study [39]. These variants were chosen based on their individual F-statistics with XL.HDL.C to be greater than 3 and across all instruments this gave a mean F-statistic of 10. The summary scatter plot for these data is shown in Figure 4. Then for the MR analyses, the summary statistics for G-X and G-Y association are extracted from 2 other independent studies [40, 34] respectively; the results for our various data analyses are given in Table 5.

**Fig. 4:**
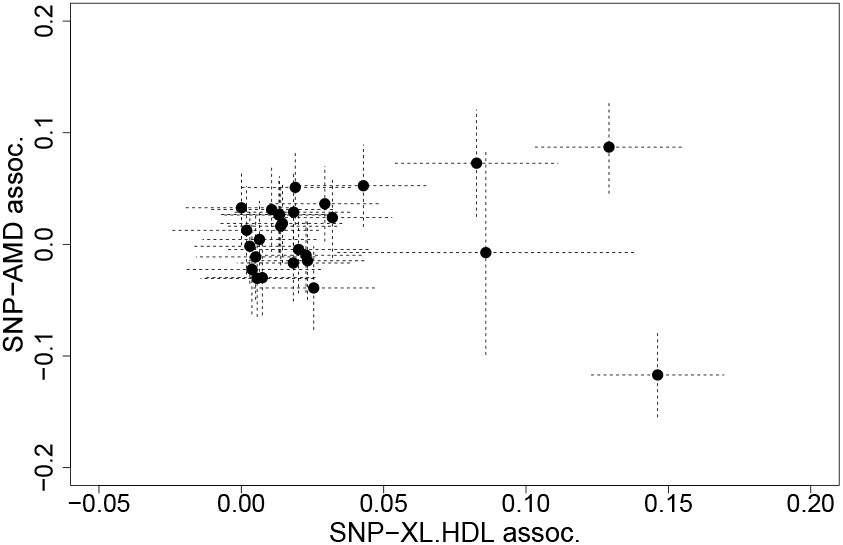
AMD and HDL: Scatter plot of the relationship between SNP-outcome and SNP-exposure association.

**Table 5:** Estimates for the causal effect of a unit increase in XL.HDL.C on the risk of AMD using a range of methods.

| Parameters | Estimator | Mean | 95% Lower Interval | 95% Upper Interval |
| --- | --- | --- | --- | --- |
| **Standard one-parameter approaches** |  |  |  |  |
| $\beta$ | IVW | 0.0251 | -0.3493 | 0.3995 |
|  | MR-APS | 0.0672 | -0.2997 | 0.4341 |
|  | MR-RAPS | 0.4567 | 0.1350 | 0.7785 |
| **BESIDE-MR: one-parameter model** |  |  |  |  |
| $\beta$ | DL estimate | 0.8331 | 0.5332 | 1.2679 |
|  | Full Bayesian | 0.8149 | 0.5050 | 1.2105 |
| $\tau^2 \times 10^{-4}$ | DL estimate | 0.0024 | 0.0000 | 0.0000 |
|  | Full Bayesian | 0.3773 | 0.0833 | 1.4330 |
| **BESIDE-MR: two-parameter model** |  |  |  |  |
| $\beta_1$ | DL estimate | 1.0219 | 0.6229 | 1.6596 |
|  | Full Bayesian | 0.9027 | 0.4998 | 1.4966 |
| $\beta_2$ | DL estimate | -0.8212 | -1.2022 | -0.4983 |
|  | Full Bayesian | -0.5948 | -1.2456 | 1.0716 |
| $\tau_1^2 \times 10^{-4}$ | DL estimate | 0.0033 | 0.0000 | 0.0000 |
|  | Full Bayesian | 0.3435 | 0.0807 | 1.2606 |
| $\tau_2^2 \times 10^{-4}$ | DL estimate | 0.0061 | 0.0000 | 0.0000 |
|  | Full Bayesian | 0.3735 | 0.0823 | 1.4568 |

When one-parameter causal models are fitted to the data, all methods estimate a positive causal association, with BESIDE-MR giving the largest effects and the IVW method giving the smallest effect. This is not surprising because the IVW estimate is known to be vulnerable to weak instrument bias towards zero in the two sample setting. Figure 5 shows the inclusion probability for each instrument, using our two implementations. The DL approach is seen to more aggressively select or de-select instruments than the full Bayesian approach.

**Fig. 5:**
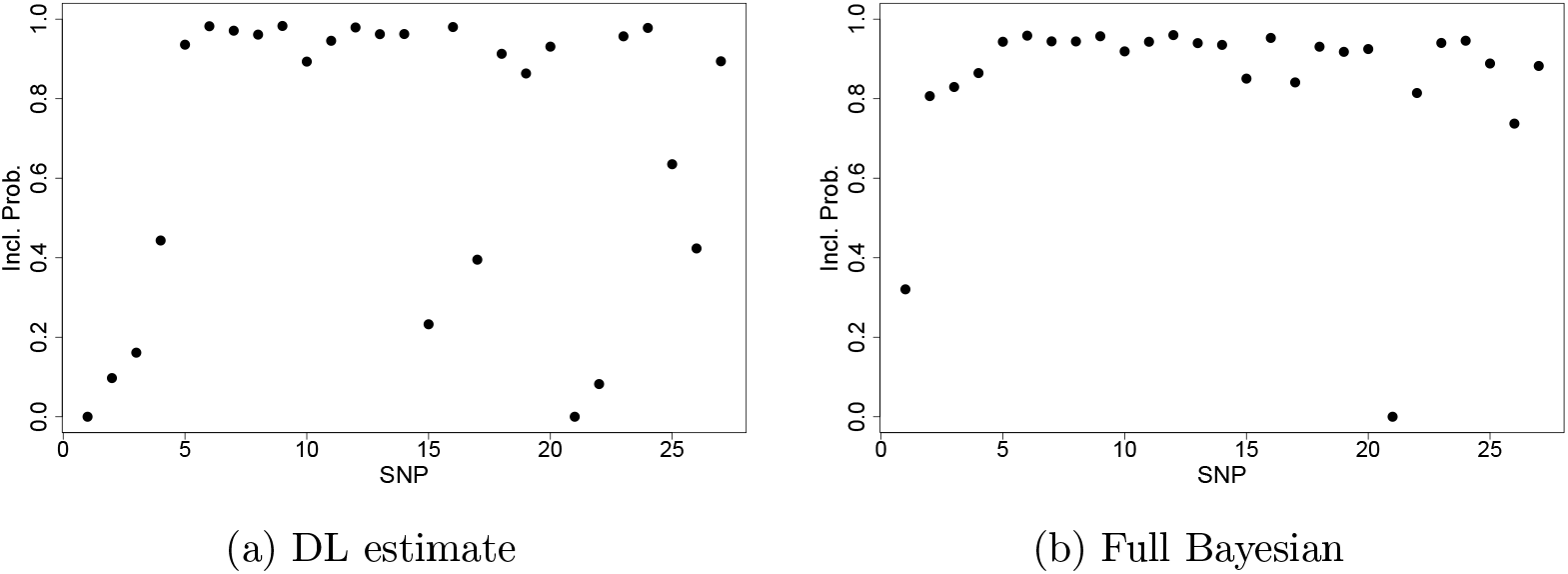
AMD and HDL: Inclusion probability (*PPI*) for each instrument for DL estimate (a) and full Bayesian approach (b).

Next, we fit our two-parameter causal model, which offers robustness to over 50% of the SNPs violating the InSIDE assumption. Interestingly, we see that this estimates two distinct causal effects of opposite sign (Table 5). For the DL approach, approximately 6 SNPs have evidence for inclusion (*PPI >* 0.75) to each of the 2 clusters, see Figure 6a. For the full Bayesian approach, 4 instruments have evidence of inclusion in the set identifying a positive relationship and only SNP *rs903319* for the negative relationship (hence 0 is within the credible interval for this smaller set), see Figure 6b. Appendix Figure A.8 show *PPI* for each instrument.

**Fig. 6:**
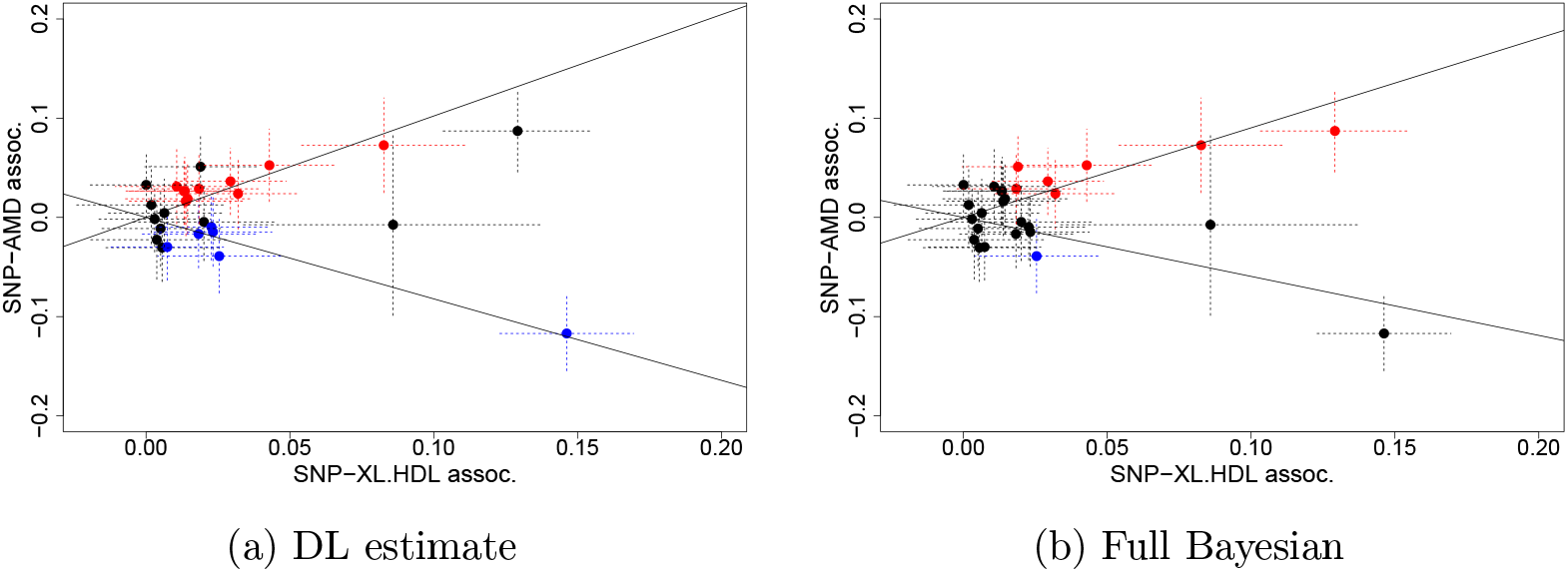
AMD and HDL: Scatter plot of the relationship between SNP-outcome and SNP-exposure association, where the coloured SNPs had *PPI >* 0.75, for DL estimate (a) and full Bayesian approach (b), assuming InSIDE violation. Colour blue and red is instrument that had strong evidence for cluster *I*_2_ that estimates *β*_2_ and *I*_1_ for *β*_1_ respectively. The solid lines are the estimated *β*_1_ and *β*_2_.

Our tentative conclusion here is that a small proportion of InSIDE-violating SNPs act to reduce the apparent causal effect of XL.HDL.C on AMD detectable by a one-parameter model. Once this set has been accounted for within a two-parameter model, this increases the evidence in favour of a causal role of XL.HDL.C on AMD further. Our results are consistent with Zuber *et al.* [38] who also found subsets of SNPs which suggested qualitatively different conclusions about the causal role of XL.HDL.C on AMD.

### 5.1 Sensitivity with penalisation for model complexity

In the simulations, the penalisation parameter for model complexity, *η* is zero, here we vary *η* between −5 to 5 for the one-parameter BESIDE-MR. Large negative *η* would force BESIDE-MR to favour models with fewer instruments and large positive *η* would favour many instruments (Table 6 and Appendix Table A.8 for *η* 2 to 5). Furthermore, shown by the heat map of *η* and *PPI*, Figure 7, the *PPI* for each instruments decreases with *η*, however there are a few instruments that have consistently higher probability for inclusion in comparison to the rest and *rs261342* is never chosen. The overall *β* did not change with *η*, but with fewer instruments BESIDE-MR becomes more uncertain of its estimation, i.e. wider credible interval. Similar patterns were found for the two parameter model, see Appendix Table A.9. The similarity in *β* estimates between ranges of *η* demonstrates that our applied example exhibits large heterogeneity and therefore only a handful of SNPs strongly influencing the results.

**Table 6:** Sensitivity analysis. Med., LCI and UCI are the median of the posterior distribution with 95% upper and lower credible intervals respectively. 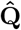 is instrument normalised Q-statistics, Σ*Q*_*j*_/*I*_*j*_. Σ*I*_*j*_ is the number of instruments included. The Q-statistic for 27 Instruments is 115.99.

| $\eta$ | Para. | Est. | -5 | | | -4 | | | -3 | | | -2 | | | -1 | | | 0 | | | 1 | | |
| --- | --- | --- | --- | --- | --- | --- | --- | --- | --- | --- | --- | --- | --- | --- | --- | --- | --- | --- | --- | --- | --- | --- | --- |
|  |  |  | Med. | LCI | UCI | Med. | LCI | UCI | Med. | LCI | UCI | Med. | LCI | UCI | Med. | LCI | UCI | Med. | LCI | UCI | Med. | LCI | UCI |
| $\beta$ | DL | | 0.78 | -1.28 | 2.02 | 0.82 | -1.21 | 2.03 | 0.90 | -0.87 | 1.96 | 0.91 | 0.39 | 1.73 | 0.85 | 0.53 | 1.49 | 0.82 | 0.53 | 1.26 | 0.80 | 0.53 | 1.12 |
|  | Bayes |  | 0.70 | -1.41 | 2.02 | 0.76 | -1.29 | 2.03 | 0.85 | -0.95 | 1.95 | 0.87 | 0.29 | 1.71 | 0.83 | 0.49 | 1.43 | 0.80 | 0.50 | 1.21 | 0.78 | 0.50 | 1.12 |
| $\tau^2 \times 10^{-4}$ | DL | | 0.00 | 0.00 | 0.00 | 0.00 | 0.00 | 0.00 | 0.00 | 0.00 | 0.00 | 0.00 | 0.00 | 0.00 | 0.00 | 0.00 | 0.00 | 0.00 | 0.00 | 0.00 | 0.00 | 0.00 | 0.00 |
|  | Bayes |  | 0.25 | 0.08 | 1.43 | 0.24 | 0.08 | 1.41 | 0.24 | 0.08 | 1.29 | 0.24 | 0.08 | 1.23 | 0.25 | 0.08 | 1.29 | 0.26 | 0.08 | 1.44 | 0.27 | 0.09 | 1.63 |
| $\hat{Q}$ | DL | | 0.56 | 0.11 | 0.98 | 0.61 | 0.12 | 0.98 | 0.75 | 0.22 | 0.99 | 0.87 | 0.46 | 1.00 | 0.93 | 0.69 | 1.00 | 0.96 | 0.80 | 1.00 | 0.98 | 0.86 | 1.00 |
|  | Bayes |  | 1.02 | 0.15 | 2.43 | 1.04 | 0.18 | 2.37 | 1.12 | 0.31 | 2.13 | 1.28 | 0.69 | 1.93 | 1.45 | 1.03 | 1.95 | 1.57 | 1.25 | 1.95 | 1.77 | 1.41 | 1.94 |
| $\sum I_j$ | DL | | 5 | 5 | 7 | 6 | 5 | 9 | 8 | 5 | 13 | 13 | 9 | 17 | 17 | 13 | 19 | 19 | 16 | 20 | 20 | 18 | 20 |
|  | Bayes |  | 5 | 5 | 7 | 6 | 5 | 9 | 9 | 5 | 13 | 15 | 10 | 19 | 20 | 15 | 23 | 23 | 20 | 25 | 25 | 22 | 26 |

**Fig. 7:**
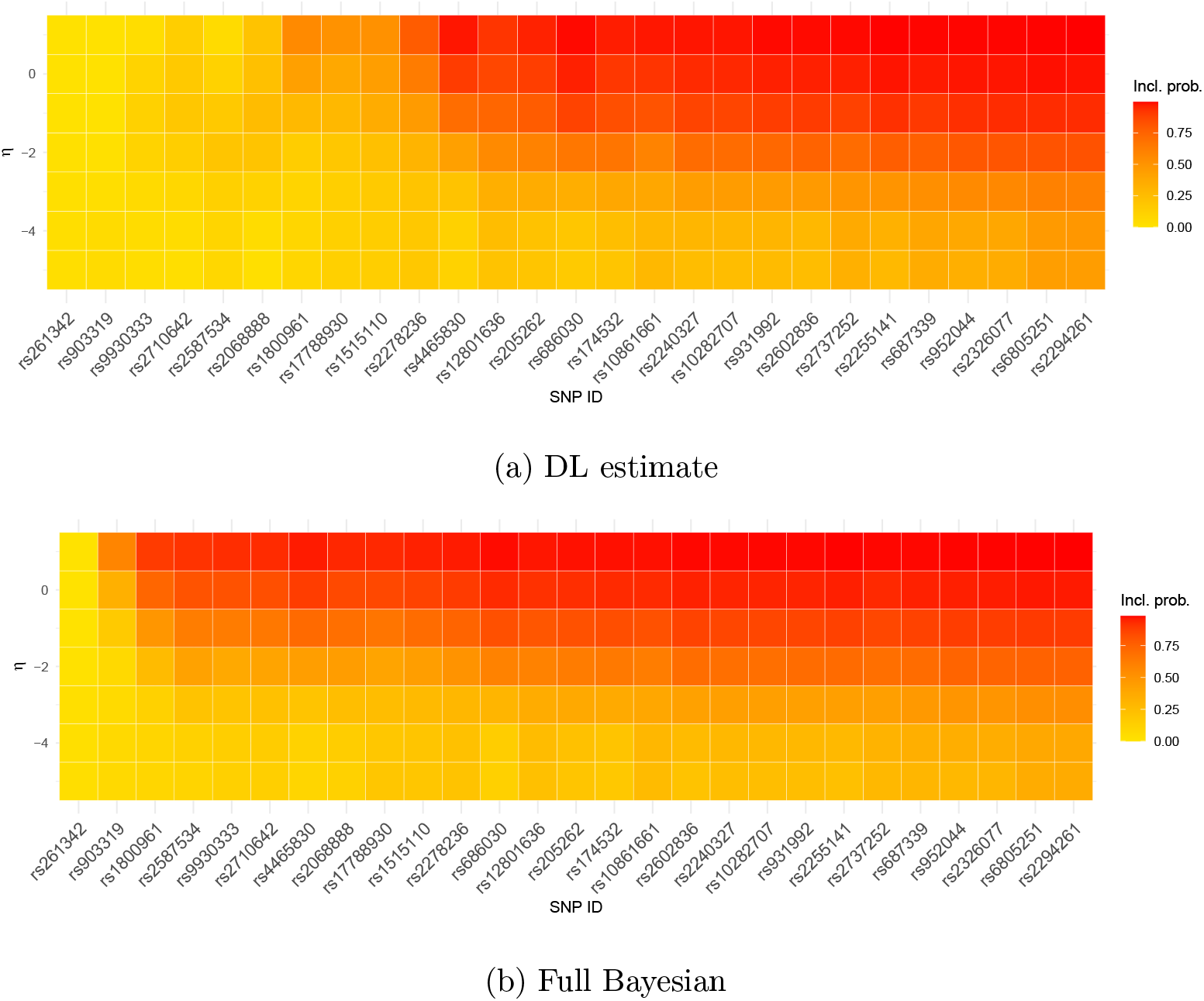
AMD and HDL *η* sensitivity: Inclusion probability for each instrument for DL estimate (a) and full Bayesian approach (b).

In the simulations, two-parameter BESIDE-MR tends focus on estimating one *β* when there is an imbalance of instruments in clusters. However, in this sensitivity analysis, BESIDE-MR consistently estimates two separate slopes over all choices of the model complexity penalization terms. This gives us confidence that the clusters are both real and robustly identified.

### 5.2 Detecting and adjusting for label switching in the full Bayesian model

The trace plots in Figure 8a and 8b show that the DL implementation consistently identifies two separate distributions for *β*_1_ and *β*_2_, which are centered around 1.02 and −0.82 respectively. This is not the case, however, under the full Bayesian implementation. Trace plots 8c and 8d show that the chains for *β*_1_ and *β*_2_ jumping between two distinct values. This is commonly known as ‘label switching’. It has been recommended that, instead of adjusting the MCMC algorithm itself, one can simply re-allocate iteration labels from the output instead [33]. To this end we performed a K-means clustering analysis [41] on the MCMC output. Before K-means correction, the mean posterior distribution of *β*_1_ and *β*_2_ gave 0.13 and 0.18 respectively. K-means analysis clustered 181,186 iterations centred at 0.90 and the second cluster contains 218,815 iterations with mean of −0.59. We re-assigned the estimates (to *β*_1_ and *β*_2_) accordingly (see Figure 8e) which gave new posterior distribution with mean and credible interval shown in Table 5. This issue further emphasizes the importance of carefully implementing the fully Bayesian approach, and for checking MCMC output for convergence issue. As an initial investigation, we added an order restriction to M-H so that *β*_1_ > *β*_2_ (we thank a reviewer for this suggestion), however, this lead to poor mixing in the MCMC run which we could not adequately address.

**Fig. 8:**
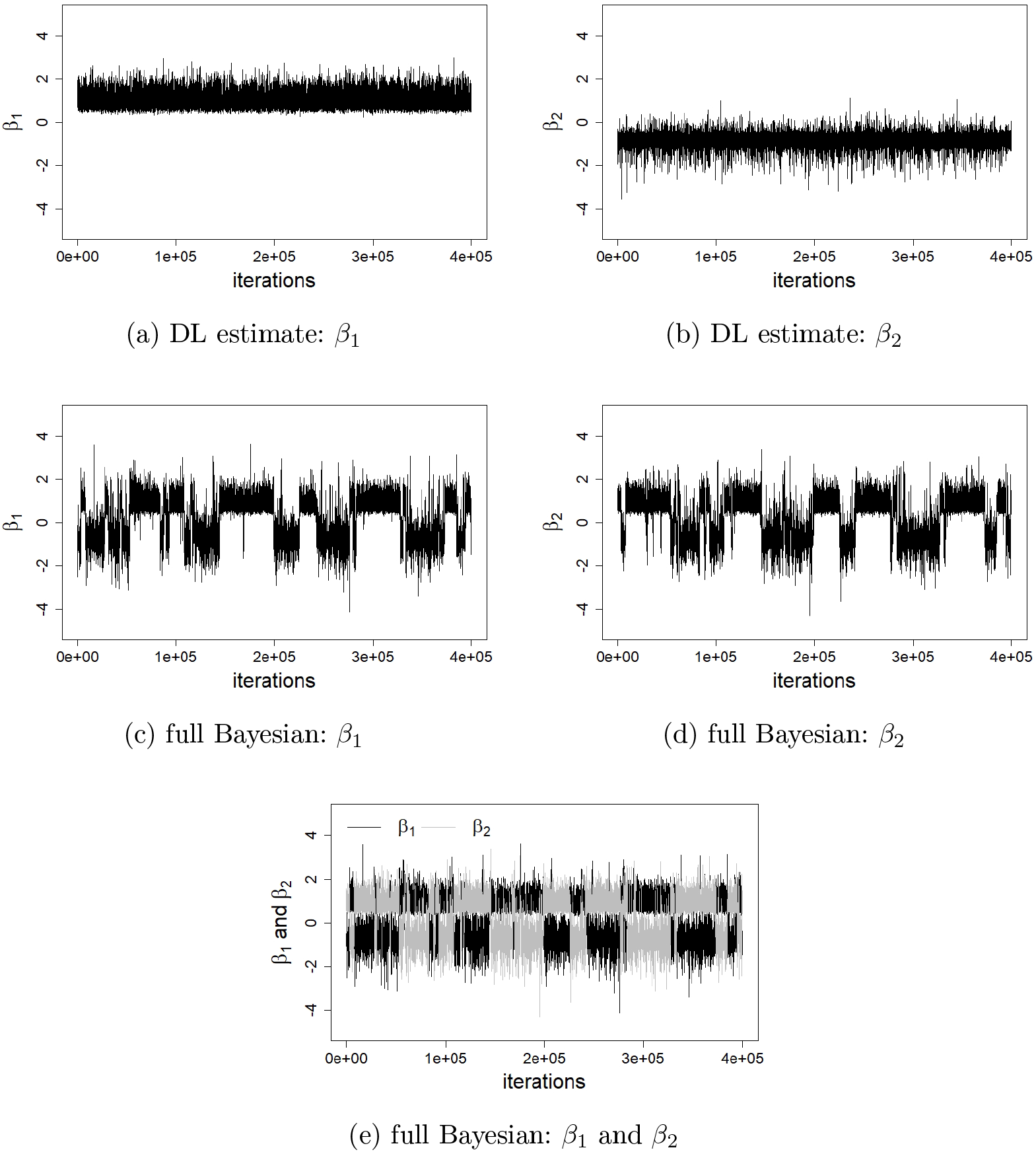
AMD and HDL: Trace plots for *β*_1_ and *β*_2_ from the full Bayesian (a,b) and DL implementations (c,d). And combined *β*_1_ and *β*_2_ for full Bayesian (e).

## 6 Discussion

In this paper we propose a Bayesian Model Averaging approach for two-sample summary data MR that offers robustness to pleiotropy and weak instruments. Our approach can be viewed as a Bayesian extension of the classical MR-RAPS approach. Rather than assuming, as MR-RAPS does, the InSIDE violating SNPs are small in number and can be effectively penalized in the analysis. Our one- and two-parameter Bayesian models go beyond this. We were able to demonstrate the potential utility of this extended model in our applied example to uncover sub-signals in the data that would be missed by conventional methods. We explored two implementations of BESIDE-MR, namely the full Bayesian and the simplified DL implementation. Our simulations showed that the DL implementation generally performed well, and led to a more aggressive selection of SNPs as either in or out of the model than the full Bayesian approach. It was also much more straightforward to fit and achieve convergence. Despite the fully Bayesian implementation requiring more computational time and careful consideration of the MCMC output, it is far better at detecting small effects and consistently identifying outlying instruments. In future work we will attempt to improve the reliability of the full Bayesian approach. Specifically, we plan to create a label switching algorithm [42] for BESIDE-MR output and specify a more sophisticated procedure for optimising the tuning parameter for each model parameter separately. In the meantime, we urge users of the full Bayesian approach to manually adapt the tuning parameters and carefully monitor the mixing and convergence of the MCMC chains, which are the essential aspects of the analysis. We also remind the reader that the number of iterations to reach convergence increases with the number of instruments. As seen in Appendix E.2, diagnostic tools such as performing multiple chains with different initial values and trace plots are useful in this regard. For a comprehensive tutorial see Albert *et al.* and Lunn *et al.* [43, 44].

A useful additional output from our BMA approach compared to classical approaches is the inclusion probability for each SNP. This of course necessitates the specification of a prior probability of inclusion, which we fixed at a constant value of 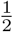. Ideally, one should use informative priors where possible. Indeed, there are multiple sources of external information, e.g. epigenetic databases and bioinformatic webtools that could be used to achieve this. For example, a genetic variant that is located in a protein coding gene relevant to the pathway between exposure and outcome of interest can be given a higher inclusion prior probability. Conversely, we might give a much lower inclusion prior probability if the variant is located in a gene that is expressed in multiple tissues. This is again a topic for future research.

It is important to note that, like other instrument selection and penalization methods, the one-parameter BESIDE-MR model assumes that the largest sets of instruments with homogeneous MR estimate consistently identify the true causal effect. However, there could be cases where this does not hold and consequently BESIDE-MR is biased. This motivated the development of our two-parameter model. The two-parameter model allows BESIDE-MR to estimate a second slope for an (approximately) equally sized instrument set of homogeneous MR estimate; as shown by the bias formulae of Equation (4), this second slope could potentially be in the form of InSIDE violation, directional pleiotropy or even represent true causal effect heterogeneity. That is, different SNPs perturb the exposure in a distinct ways that gives rise to two true causal effects. This possibility is explored in recent work by Iong *et al.* [45]. In future work we plan to explore the utility of BESIDE-MR in this alternative setting as well.

Zuber *et al.* [38] have proposed a BMA implementation of multivariable MR [37, 46]. Our model can in principle be extended to multivariable MR too. For a model with 10 exposure traits, this would necessitate the estimation of 11 causal parameters to account for InSIDE violation via unmeasured pathways. This is a potential topic for future research. BESIDE-MR could also be extended to correlated SNPs and 2 dependent samples, both will require additional weights to account for correlation between SNPs for the former, and correlations between the SNP-exposure and SNP-outcome association estimates for the latter.

## Software

Software in the form of R code is available on corresponding author’s `Github` (https://github.com/CYShapland/BESIDEMR).

## Acknowledgements

We thank the reviewers and associate editor for providing comments and suggestions to improve this paper. C.Y.S. and J.B. works in a unit that receives support from the University of Bristol and the UK Medical Research Council.

## A Assumptions for two-sample MR analyses

Table A.1 gives the summary of the assumptions made, which closely follows Table 1 in Bowden *et al.*[7] with exception to NO Measurement Error (NOME) assumption, as the measurement error is computed in our profile likelihood. For the estimation of local average causal effect, additional *structural* assumption is required, that is the model is linear and additive without interactions. The structural assumption could be violated in number of situations [47], in most MR applications scenarios of, binary outcomes and interaction between X and G, is plausible. The former we will discuss in Section 7, and for violation from the latter, approximation of local average causal effect using the linear structural model will still hold in many cases as most of the SNPs effect on X is usually very small [31]. Variation in Instrument Strength (VIS) is reasonable as we assume some sampling error would exist and SNPs used are uncorrelated.

**Table A.1:** Summary of assumptions for two-sample MR analyses. G is the genetic instrument, X is the exposure, Y is the outcome and Z is the unmeasured confounding.

| Assumptions | Description |
| --- | --- |
| **Instrumental variable (IV) specific** |  |
| IV1 | G is associated with X, not through U ($\gamma_j > 0$). |
| IV2 | G is not associated with U ($\psi_j = 0$). |
| IV3 | No direct effect between G and Y ($\alpha_j = 0$). |
| Structural | Model (2.1) and (2.2) are linear and without interaction. |
| **Two-sample Analyses (TSA) specific** |  |
| TSA1 | Model (2.1) and (2.2) holds for sample 1 and 2. |
| TSA2 | $\epsilon_k^Y$ and $\epsilon_k^{Y'}$ in Model (2.1) and (2.2) respectively are independent from each other and other variables. |
| TSA3 | $\sigma_{X_j}^2$ and $\sigma_{Y_j}^2$ in Model (2.3) are independent and known. |
| **Instrument Strength Independent of Direct Effect (InSIDE), under IV2** |  |
| General InSIDE | $\widehat{Cov}(\alpha_j, \gamma_j) \rightarrow 0$ as $L \rightarrow \infty$. |
| Perfect InSIDE | $\widehat{Cov}(\alpha_j, \gamma_j) = 0$ for analysed sample. |
| **Variation in Instrument Strength (VIS), under IV2** |  |
| VIS | $\gamma_j \neq \gamma_i$ from $G_i$ and $G_j$ on X. |
| **One parameter BESIDE-MR** |  |
| ZERO Modal Pleiotropy Assumption (ZEMPA) | largest set of instruments with homogeneous causal effect estimates. |

## B Bias from violation of InSIDE assumption

Suppose we are estimating the causal parameter from instruments that violate the InSIDE assumption using the IVW approach. Its estimand will equal:

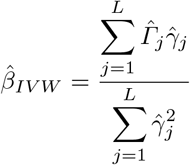

as the 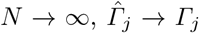 and 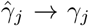, so that asymptotically, the expectation of IVW estimate tends towards the following

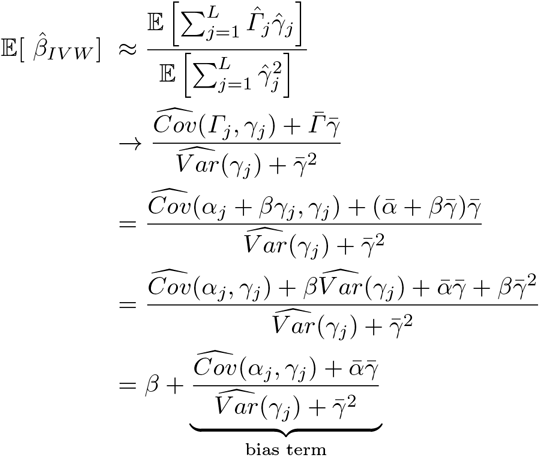

When InSIDE is perfectly violated (*α*_*j*_=*γ*_*j*_) the numerator and denominator of the bias term are equal. Therefore, *β*_*IVW*_ =*β*+1

## C Metropolis-Hastings algorithm for the one-parameter causal model

For updating the model parameter values, instead of using the standard Gibbs sampling, where it requires conditional posterior distribution, we used a random walk M-H algorithm to give a proposal distribution for each parameter. Let 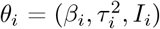 be the current *i*th value of the parameter vector *θ*. *θ*_*i*_ is updated to *θ*_*i*+1_ one parameter at a time, by simulating a candidate value *θ*\* from proposal density, until it is accepted. Note that if the proposal density *C*() for a given parameter is ‘symmetric’ - that is if *C*(*θ*_*i*_|*θ*_*i*+1_) = *C*(*θ*;_*i*+1_|*θ*_*i*_) then the proposal density can be omitted from the calculation of the acceptance probability. This is the case for *β* and *I*, but not *τ*^2^.

### C.1 The full Bayesian implementation

**– Update** *β*

1. Sample *β*\* ~ *β*_*i*_ + *h*_*β*_*N*(0, 1), where *h*_*β*_ is a user defined tuning constant.
2. Accept *β*_*i*+1_ = *β*\* with probability:

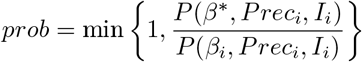

otherwise set *β*_*i*+1_ = *β*_*i*_, where *P*(,) is the posterior density.
– **Update** *Prec* (*τ*^2^ = 1/*Prec*)

1. Sample

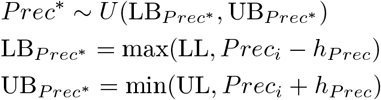

where *U*(,) is the proposal density in the form of a uniform distribution. LL and UL is user defined lower and upper limit for *Prec* respectively, and *h*_*Prec*_ is a user defined tuning constant.
2. Accept *Prec*_*i*+1_ = *Prec*\* with probability:

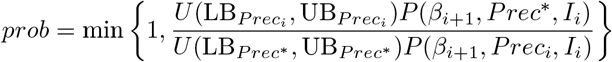 Otherwise set *Prec*_*i*+1_ = *Prec*_*i*_, where *P*(,) is the posterior density.
– **Update** *I*

1. Generate a random number between 1 and *L* from *P*(*I*_*L*_), define it as 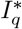, which is the *q*th element of *I*\*.
2. Set 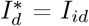 for all *d* ≠ *q*.
3. Set 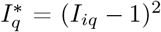 (this defines the proposed and current model to differ by one instrument).
4. If 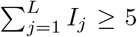 is true, continue to the next step, otherwise repeat step 1 (ensures there is enough IVs for estimation).
5. Accept *I*_*i*+1_ = *I*\* with probability:

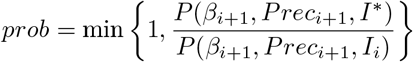

otherwise set *I*_*i*+1_ = *I*_*i*_,, where *P*(,) is the posterior density.

The *h*_*β*_ and *h*_*Prec*_ acts as tuning parameters for the acceptance rate. That is, the proportion of iterations that *θ*\* is accepted as *θ*_*i*+1_. Acceptance rates are recommended to be between 0.25 and 0.45 for the random walk M-H algorithm [43]. We follow this guidance in our implementation of the approach in simulations and applied data settings.

### C.2 The DL implementation

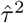 is calculated from DerSimonian-Laird estimate [30] and estimated from every proposed value of *β* and *L*;

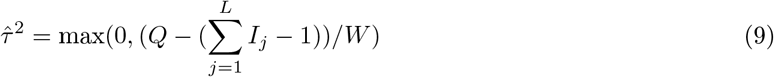

where

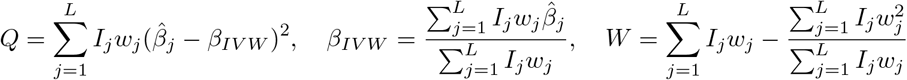

and 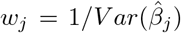 respectively. Note that *I*_*j*_ should not be confused with Higgin’s *I*^2^ statistic used to quantify heterogeneity in meta-analysis.

– **Update** *β*

1. Smaple *β*\* ~ *β*_*i*_+*h*_*β*_*N*(0, 1) where *h*_*β*_ is a user defined tuning constant.
2. Accept *β*_*i*+1_ = *β*\* with probability:

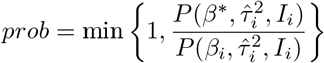 Otherwise set *β*_*i*+1_ = *β*_*i*_, *P*(,) is the posterior density.
– **Update** *L*

1. Generate a random number between 1 and *L* from *P*(*I*_*L*_), define it as 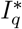, which is the *q*th element of *I*\*.
2. Set 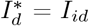 for all *d* ≠ *q*.
3. Set 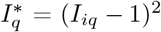.
4. If 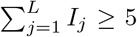 is true, continue to the next step, otherwise repeat step 1 (ensures there is enough IVs for estimation).
5. Accept *I*_*i*+1_ = *I*\* with probability:

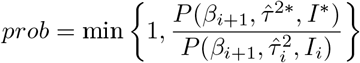

where 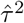 and 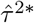 is calculated with *I*_*i*_ and *I*\* respectively. Otherwise set *I*_*i*+1_ = *I*_*i*_, where *P*(,) is the posterior density.

## D Derivation of integrated likelihood

Based on the model shown in Equation (3), and the instruments included and excluded have:

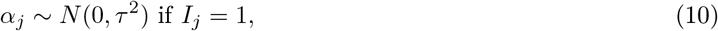

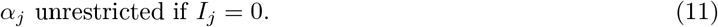

then the likelihood function for summary data of the *G* − *X* and *G* − *Y* can be given by as:

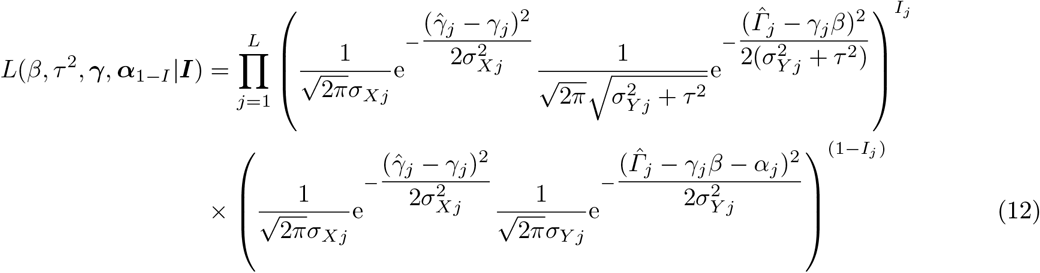

The integrated likelihood of *β* and *τ*^2^ is then defined as:

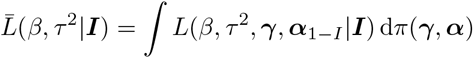

for some distribution on (***γ***, ***α***). We can approximate the integrated likelihood by using Laplace method:

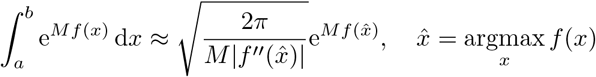

Let ***θ*** = (***γ***, ***α***_1−*I*_) and assume it is flat, then

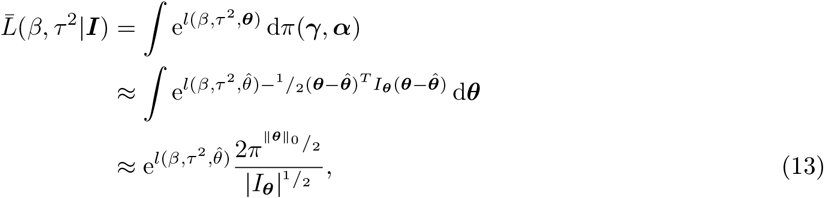

where 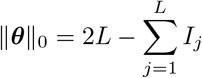 and *I*_***θ***_ = d*iag*(*I*_***θ***1_, · · ·, *I*_***θ**L*_).

We can profile out 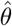 from 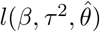 to give the profile likelihood of (*β, τ*^2^):

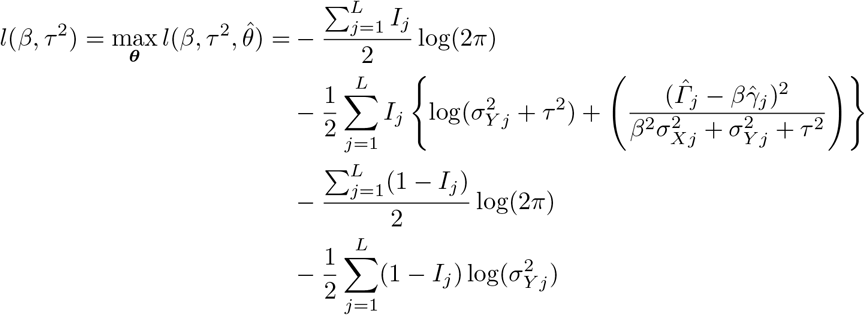

Then our integrated likelihood is:

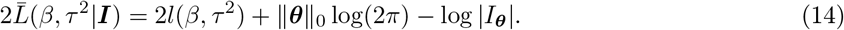

*I*_***θ***_ is the Fisher information matrix for

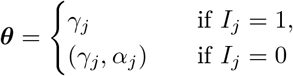

so that

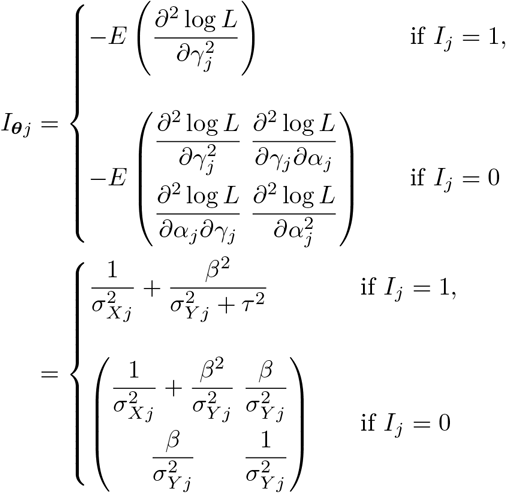

therefore the sum of the log determinant of the information matrix is:

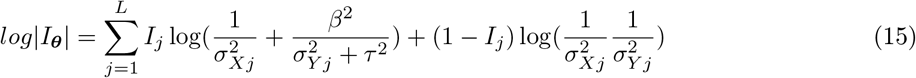

if *β* ≈ 0, then

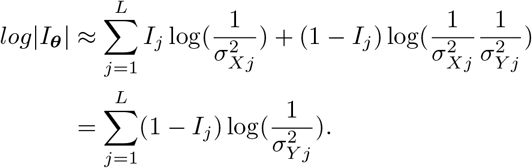

With this, Equation 14 approximates to,

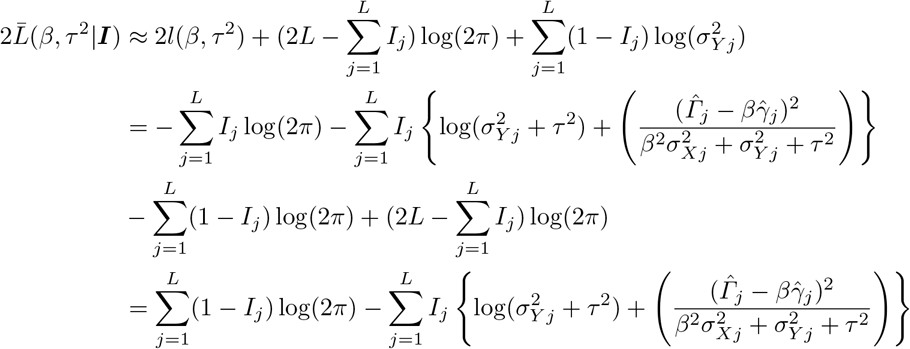

## E Simulations under the one-parameter model

This section is specifically for one-parameter BESIDE-MR, that covers Monte Carlo simulation method, and results for convergence, weaker instruments (L=50 and mean F-statistics of 10), many weak instruments (L=100, mean F-statistics of 5 and 10) and sensitivity to strengths of heterogeneity (varied Q-statistics).

### E.1 Simulation Method

We simulate two-sample summary MR data sets with *L*=50 instruments from model;

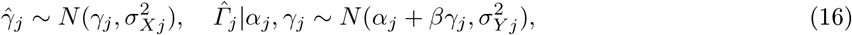

The parameters *γ*_*j*_ were generated from a Uniform *U*(0.34, 1.1) distribution, *σ*_*Xj*_ was generated from a Uniform *U*(0.06, *UB*) and *σ*_*Yj*_ was generated from a Uniform *U*(0.015, 0.11) distribution. The upper bound on the G-X association standard error *UB* was used to determine mean instrument strength - with 0.095 ≤ *UB* ≤ 1 giving mean F-statistics between 10 and 100 respectively. In this setting, the F-statistic for a single SNP can be approximated as 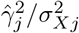. We defined invalid instruments as SNPs that have non-zero *α*_*j*_, as there is a direct effect from SNP to outcome, i.e. violation to IV3.

*α*_*j*_ for invalid instruments is simulated from normal *N*(*μ*_*α*_, 0.04) distribution, with the parameter *μ*_*α*_ being used to determine the mean bias induced by including the invalid instruments in the model. The task of BESIDE-MR in the presence of a non-zero *μ*_*α*_ is to give large weight to models which include SNPs for which *μ*_*α*_ ≈ 0. As summarised by Table 1 in the main manuscript, *μ*_*α*_ = 0 for the instruments that have balanced pleiotropic effect, and *μ*_*α*_ = 0.05 for directional pleiotropic effect. Apart from a potential non-zero mean bias, the simulated pleiotropic effects satisfy the InSIDE assumption.

For evaluation criteria, we monitor the following quantities across our simulations:

– Mean bias of the causal parameter estimate. For BESIDE-MR we use the mean of the posterior distribution of *β* to assess this;
– Coverage: For IVW, MR-APS and MR-RAPS this is based on 95% symmetric confidence intervals assuming normality. For BESIDE-MR this is based on a 95% credibility interval;
– The difference in inclusion probability between valid and invalid SNPs set (BESIDE-MR only): 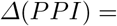 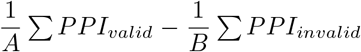, where *A* and *B* is total number of valid and invalid instruments respectively, and hence *A* + *B* = *L*.

We also report the exact *Q*-statistic [8]:

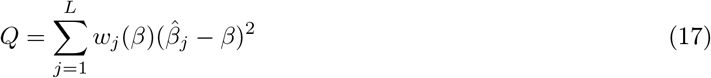

where 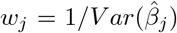. Note that only invalid SNPs which have a non-zero pleiotropic effect make a non-nominal contribution, so that, for a fixed set of pleiotropy parameters *α*_1_, *..., α*_*L*_:

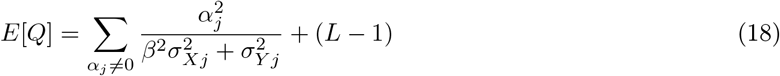

from knowing that [8];

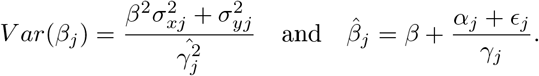

### E.2 Convergence

Convergence is an important aspect to Bayesian analysis when implemented using MCMC methods, as it is an iterative process, different possible values are explored at each iteration. To investigate convergence, we run 5 short chains, each with random starting values, 50,000 iterations and 10,000 burn-ins. We also run one long chain with 500,000 iterations and 100,000 burn-ins.

We tested convergence on 3 different types of instruments; (1) Scenario 1 and (2) Scenario 2 without invalid instruments, and (3) Scenario 1 with 30% invalid instruments.

Table A.2 demonstrates evidence for convergence with 50,000 iterations and 10,000 burn in. The mean, standard deviation and 95% credible interval of the posterior distribution for *β* are similar between long and shorts chains, in all 3 scenarios. The difference shown between long and short chains are the standard error and the time-series standard error (adjusted for auto-correlation). This is expected as the accuracy for the posterior mean of *β* increases with number of iterations. The trace plot is another diagnostic tool; it is a continuous line that shows the values a parameter has against the iteration number. A “catepillar” shaped trace plot, and similarities between long and short chains, supports evidence for convergence (Figure A.1 and A.2). Table A.3 gives the *PPI* of the 10 SNPs from long and short chain. These 10 SNPs were selected because they had the highest *PPI* in the long chain. The similarity in inclusion probability between the short and long chains for all the 10 instruments and across scenarios (Table A.3) demonstrates evidence for convergence in *PPI*.

**Fig. A.1:**
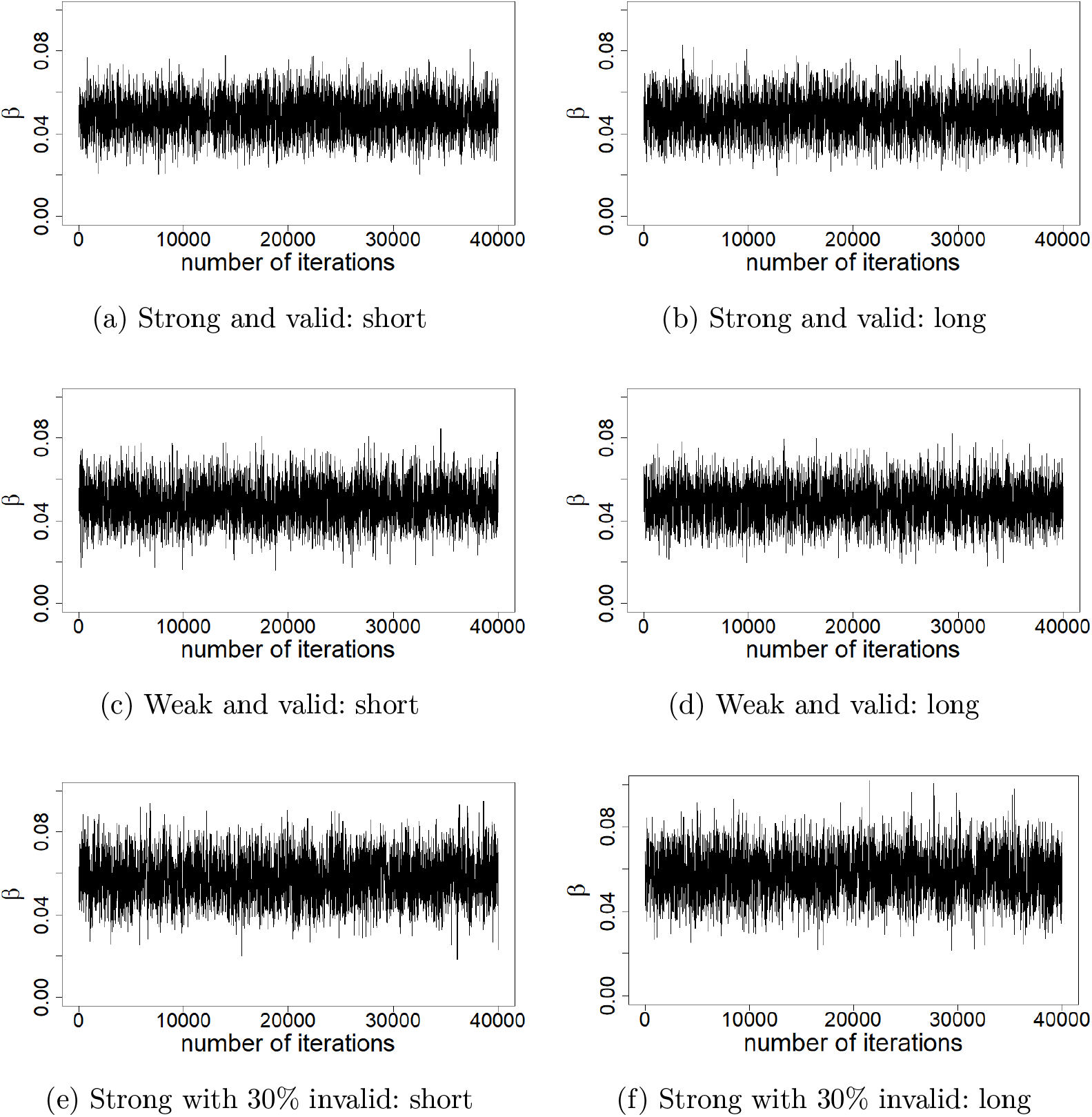
Trace plot of the causal effect estimate (*β*) from DL approach with 3 different instrument scenarios; (a, b) strong valid, (c, d) weak valid instruments only and (e, f) strong with 30% invalid instruments. Short and long chain consist of 50,000 and 500,000 iterations with 10,000 and 100,000 burn-in respectively.

**Table A.2:** Convergence diagnostic of Scenario 1 and 2 without invalid, and with 30% invalid instruments by comparing a long and 5 short chains. Each short chain have 50,000 iterations with 5,000 burn-ins and the long chain have 500,000 iterations and 100,000 burn-ins. True *β* is 0.05. SD, standard deviation; SE, standard error; CI, credible interval; inst., instrument(s).

| Inst.<br>scenario | chain | mean $\beta$ | SD | SE | Time-series<br>SE | Lower<br>95% CI | Upper<br>95% CI |
| --- | --- | --- | --- | --- | --- | --- | --- |
| Strong<br>and valid | **DL estimate** |  |  |  |  |  |  |
|  | 1 | 0.0485 | 0.0080 | 0.00004 | 0.00014 | 0.0332 | 0.0640 |
|  | 2 | 0.0488 | 0.0081 | 0.00004 | 0.00014 | 0.0330 | 0.0643 |
|  | 3 | 0.0485 | 0.0079 | 0.00004 | 0.00014 | 0.0333 | 0.0643 |
|  | 4 | 0.0484 | 0.0081 | 0.00004 | 0.00014 | 0.0324 | 0.0645 |
|  | 5 | 0.0485 | 0.0081 | 0.00004 | 0.00014 | 0.0325 | 0.0642 |
|  | Long | 0.0485 | 0.0081 | 0.00001 | 0.00005 | 0.0327 | 0.0645 |
|  | **Full Bayesian** |  |  |  |  |  |  |
|  | 1 | 0.0486 | 0.0082 | 0.00004 | 0.00014 | 0.0323 | 0.0647 |
|  | 2 | 0.0489 | 0.0080 | 0.00004 | 0.00013 | 0.0331 | 0.0646 |
|  | 3 | 0.0487 | 0.0082 | 0.00004 | 0.00014 | 0.0324 | 0.0647 |
|  | 4 | 0.0487 | 0.0083 | 0.00004 | 0.00013 | 0.0327 | 0.0650 |
|  | 5 | 0.0491 | 0.0083 | 0.00004 | 0.00015 | 0.0325 | 0.0653 |
|  | Long | 0.0488 | 0.0083 | 0.00001 | 0.00005 | 0.0326 | 0.0652 |
| Weak<br>and valid | **DL estimate** |  |  |  |  |  |  |
|  | 1 | 0.0485 | 0.0083 | 0.00004 | 0.00016 | 0.0328 | 0.0652 |
|  | 2 | 0.0482 | 0.0083 | 0.00004 | 0.00015 | 0.0323 | 0.0648 |
|  | 3 | 0.0480 | 0.0083 | 0.00004 | 0.00015 | 0.0324 | 0.0649 |
|  | 4 | 0.0484 | 0.0083 | 0.00004 | 0.00014 | 0.0329 | 0.0652 |
|  | 5 | 0.0484 | 0.0083 | 0.00004 | 0.00016 | 0.0329 | 0.0648 |
|  | Long | 0.0482 | 0.0083 | 0.00001 | 0.00005 | 0.0323 | 0.0650 |
|  | **Full Bayesian** |  |  |  |  |  |  |
|  | 1 | 0.0483 | 0.0084 | 0.00004 | 0.00014 | 0.0319 | 0.0653 |
|  | 2 | 0.0481 | 0.0085 | 0.00004 | 0.00015 | 0.0320 | 0.0656 |
|  | 3 | 0.0485 | 0.0083 | 0.00004 | 0.00015 | 0.0324 | 0.0652 |
|  | 4 | 0.0485 | 0.0084 | 0.00004 | 0.00016 | 0.0330 | 0.0657 |
|  | 5 | 0.0480 | 0.0086 | 0.00004 | 0.00015 | 0.0318 | 0.0655 |
|  | Long | 0.0484 | 0.0085 | 0.00001 | 0.00005 | 0.0322 | 0.0658 |
| Strong<br>with<br>30%<br>invalid | **DL estimate** |  |  |  |  |  |  |
|  | 1 | 0.0578 | 0.0094 | 0.00005 | 0.00020 | 0.0397 | 0.0768 |
|  | 2 | 0.0580 | 0.0097 | 0.00005 | 0.00024 | 0.0396 | 0.0776 |
|  | 3 | 0.0575 | 0.0094 | 0.00005 | 0.00020 | 0.0389 | 0.0767 |
|  | 4 | 0.0573 | 0.0094 | 0.00005 | 0.00024 | 0.0395 | 0.0767 |
|  | 5 | 0.0584 | 0.0098 | 0.00005 | 0.00026 | 0.0401 | 0.0789 |
|  | Long | 0.0576 | 0.0095 | 0.00002 | 0.00008 | 0.0391 | 0.0766 |
|  | **Full Bayesian** |  |  |  |  |  |  |
|  | 1 | 0.0574 | 0.0097 | 0.00005 | 0.00023 | 0.0384 | 0.0768 |
|  | 2 | 0.0580 | 0.0096 | 0.00005 | 0.00022 | 0.0394 | 0.0769 |
|  | 3 | 0.0574 | 0.0097 | 0.00005 | 0.00021 | 0.0385 | 0.0767 |
|  | 4 | 0.0576 | 0.0096 | 0.00005 | 0.00023 | 0.0394 | 0.0768 |
|  | 5 | 0.0574 | 0.0096 | 0.00005 | 0.00021 | 0.0384 | 0.0765 |
|  | Long | 0.0576 | 0.0095 | 0.00001 | 0.00007 | 0.0392 | 0.0764 |

**Table A.3:** *PPI* of the 10 SNPs from short and long chains to diagnose the convergence of instrument probability. Note that the SNPs shown are the ones with the highest *PPI* in the long chain and for each scenario these SNPs differs. Each short chain have 50,000 iterations with 5,000 burn-ins and the long chain have 500,000 iterations and 100,000 burn-ins. inst., instrument(s).

| Inst.<br>scenario | chain | SNP1 | SNP2 | SNP3 | SNP4 | SNP5 | SNP6 | SNP7 | SNP8 | SNP9 | SNP10 |
| --- | --- | --- | --- | --- | --- | --- | --- | --- | --- | --- | --- |
| Strong<br>and valid | **DL estimate** |  |  |  |  |  |  |  |  |  |  |
|  | 1 | 0.96 | 0.97 | 0.95 | 0.93 | 0.91 | 0.92 | 0.93 | 0.90 | 0.91 | 0.92 |
|  | 2 | 0.98 | 0.96 | 0.93 | 0.92 | 0.92 | 0.92 | 0.93 | 0.93 | 0.91 | 0.93 |
|  | 3 | 0.97 | 0.96 | 0.94 | 0.94 | 0.92 | 0.92 | 0.92 | 0.93 | 0.91 | 0.91 |
|  | 4 | 0.97 | 0.95 | 0.92 | 0.92 | 0.92 | 0.91 | 0.93 | 0.92 | 0.92 | 0.90 |
|  | 5 | 0.97 | 0.95 | 0.94 | 0.94 | 0.93 | 0.91 | 0.94 | 0.92 | 0.91 | 0.88 |
|  | Long | 0.96 | 0.96 | 0.94 | 0.93 | 0.92 | 0.91 | 0.92 | 0.92 | 0.92 | 0.92 |
|  | **Full Bayesian** |  |  |  |  |  |  |  |  |  |  |
|  | 1 | 0.96 | 0.96 | 0.93 | 0.92 | 0.92 | 0.92 | 0.93 | 0.91 | 0.93 | 0.91 |
|  | 2 | 0.95 | 0.96 | 0.94 | 0.94 | 0.91 | 0.91 | 0.91 | 0.93 | 0.92 | 0.91 |
|  | 3 | 0.97 | 0.95 | 0.93 | 0.94 | 0.93 | 0.92 | 0.92 | 0.93 | 0.90 | 0.91 |
|  | 4 | 0.94 | 0.96 | 0.95 | 0.91 | 0.91 | 0.90 | 0.90 | 0.88 | 0.93 | 0.92 |
|  | 5 | 0.96 | 0.95 | 0.94 | 0.91 | 0.93 | 0.92 | 0.94 | 0.92 | 0.91 | 0.91 |
|  | Long | 0.96 | 0.96 | 0.94 | 0.93 | 0.92 | 0.92 | 0.92 | 0.92 | 0.92 | 0.91 |
| Weak<br>and valid | **DL estimate** |  |  |  |  |  |  |  |  |  |  |
|  | 1 | 0.96 | 0.95 | 0.94 | 0.92 | 0.90 | 0.91 | 0.92 | 0.90 | 0.90 | 0.89 |
|  | 2 | 0.94 | 0.96 | 0.96 | 0.93 | 0.92 | 0.92 | 0.92 | 0.91 | 0.93 | 0.90 |
|  | 3 | 0.96 | 0.97 | 0.94 | 0.95 | 0.92 | 0.93 | 0.91 | 0.92 | 0.92 | 0.91 |
|  | 4 | 0.96 | 0.97 | 0.92 | 0.93 | 0.92 | 0.92 | 0.92 | 0.92 | 0.91 | 0.90 |
|  | 5 | 0.95 | 0.96 | 0.94 | 0.95 | 0.93 | 0.91 | 0.92 | 0.90 | 0.92 | 0.93 |
|  | Long | 0.96 | 0.95 | 0.94 | 0.94 | 0.91 | 0.92 | 0.91 | 0.92 | 0.92 | 0.91 |
|  | **Full Bayesian** |  |  |  |  |  |  |  |  |  |  |
|  | 1 | 0.96 | 0.97 | 0.95 | 0.92 | 0.90 | 0.91 | 0.90 | 0.91 | 0.90 | 0.93 |
|  | 2 | 0.94 | 0.97 | 0.94 | 0.93 | 0.92 | 0.92 | 0.92 | 0.93 | 0.90 | 0.93 |
|  | 3 | 0.96 | 0.96 | 0.94 | 0.95 | 0.91 | 0.89 | 0.92 | 0.90 | 0.92 | 0.92 |
|  | 4 | 0.95 | 0.97 | 0.94 | 0.93 | 0.90 | 0.90 | 0.90 | 0.91 | 0.90 | 0.88 |
|  | 5 | 0.95 | 0.96 | 0.94 | 0.94 | 0.94 | 0.92 | 0.92 | 0.95 | 0.91 | 0.93 |
|  | Long | 0.96 | 0.96 | 0.94 | 0.93 | 0.92 | 0.92 | 0.92 | 0.91 | 0.91 | 0.91 |
| Strong<br>with<br>30%<br>invalid | **DL estimate** |  |  |  |  |  |  |  |  |  |  |
|  | 1 | 0.96 | 0.97 | 0.95 | 0.92 | 0.94 | 0.93 | 0.92 | 0.88 | 0.90 | 0.88 |
|  | 2 | 0.94 | 0.96 | 0.93 | 0.93 | 0.93 | 0.93 | 0.93 | 0.91 | 0.90 | 0.91 |
|  | 3 | 0.97 | 0.96 | 0.93 | 0.92 | 0.96 | 0.93 | 0.94 | 0.92 | 0.93 | 0.90 |
|  | 4 | 0.94 | 0.95 | 0.93 | 0.94 | 0.94 | 0.94 | 0.93 | 0.91 | 0.93 | 0.90 |
|  | 5 | 0.96 | 0.96 | 0.93 | 0.93 | 0.94 | 0.94 | 0.95 | 0.89 | 0.91 | 0.89 |
|  | Long | 0.95 | 0.95 | 0.94 | 0.94 | 0.93 | 0.93 | 0.93 | 0.92 | 0.92 | 0.90 |
|  | **Full Bayesian** |  |  |  |  |  |  |  |  |  |  |
|  | 1 | 0.94 | 0.94 | 0.94 | 0.94 | 0.94 | 0.93 | 0.95 | 0.92 | 0.93 | 0.90 |
|  | 2 | 0.94 | 0.95 | 0.93 | 0.92 | 0.95 | 0.95 | 0.92 | 0.93 | 0.87 | 0.91 |
|  | 3 | 0.95 | 0.95 | 0.93 | 0.94 | 0.95 | 0.92 | 0.93 | 0.91 | 0.93 | 0.90 |
|  | 4 | 0.97 | 0.94 | 0.94 | 0.93 | 0.92 | 0.93 | 0.90 | 0.90 | 0.89 | 0.91 |
|  | 5 | 0.94 | 0.96 | 0.94 | 0.96 | 0.94 | 0.94 | 0.91 | 0.92 | 0.92 | 0.88 |
|  | Long | 0.96 | 0.96 | 0.93 | 0.93 | 0.93 | 0.93 | 0.93 | 0.92 | 0.91 | 0.90 |

**Fig. A.2:**
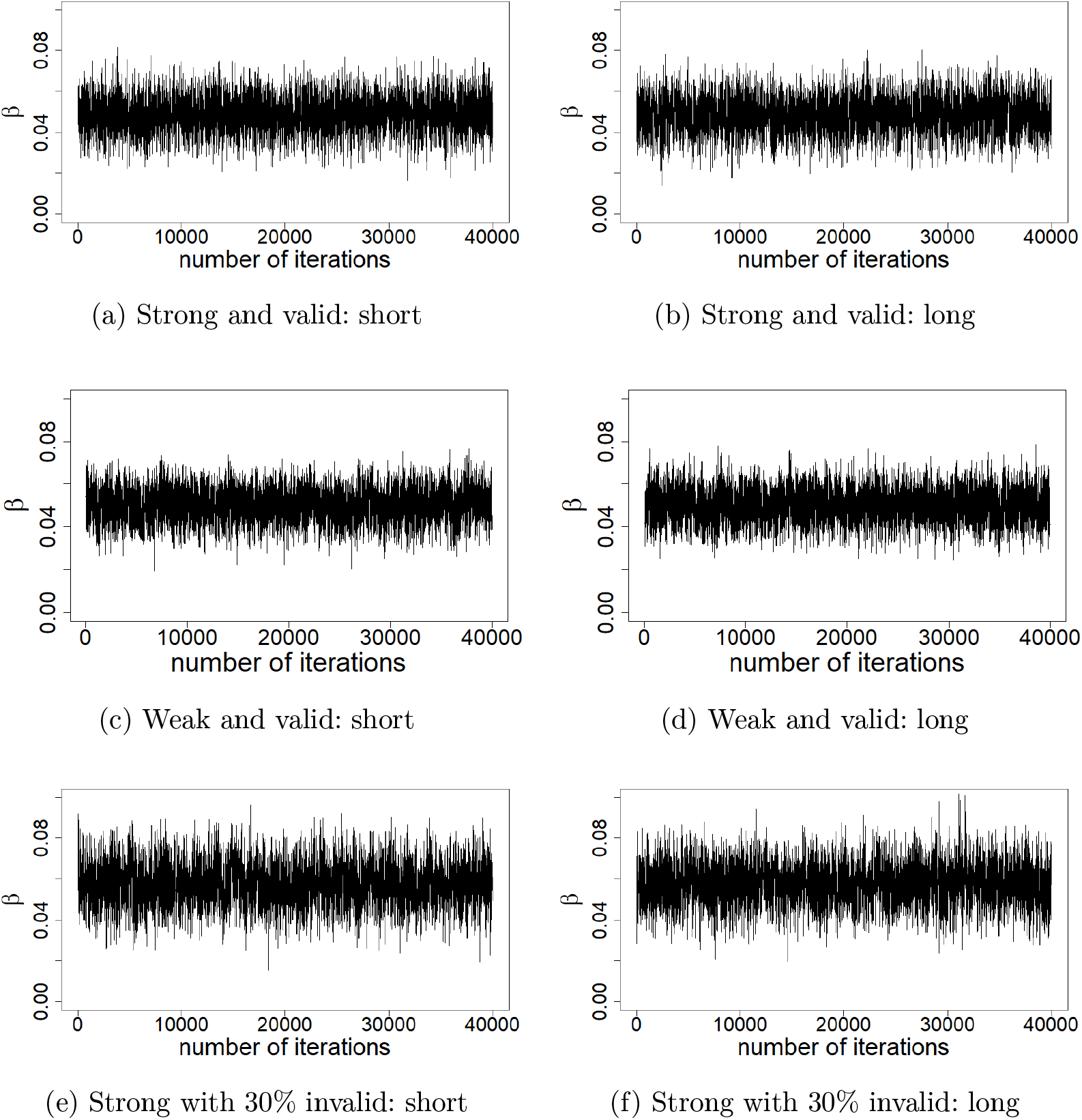
Trace plot of the causal effect estimate (*β*) from full Bayesian approach with 3 different instrument scenarios; (a, b) strong valid, (c, d) weak valid instruments only and (e, f) strong with 30% invalid instruments. Short and long chain consist of 50,000 and 500,000 iterations with 10,000 and 100,000 burn-in respectively.

### E.3 Weaker instruments

**Fig. A.3:**
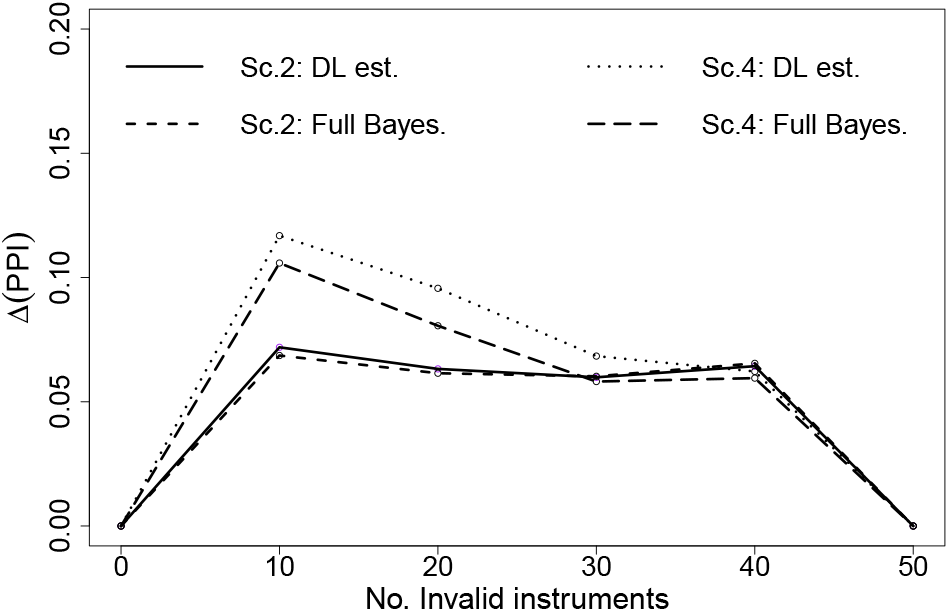
The difference in posterior probabilities of inclusion (*PPI*) between valid and invalid instruments for balanced and directional pleiotropy (Scenario 2 and 4 respectively). On the x-axis is the number of invalid/pleiotropic instruments, and the y-axis is the average difference in PPI in valid and invalid instruments set, *Δ*(*PPI*), over 1,000 simulations. As shown by legend within plot, the lines denotes results from different implementation of BESIDE-MR within each scenario.

### E.4 Many weak instruments

Many weak instruments were simulated under scenario 1, but with 100 instruments. We experimented with 2 different mean F-statistics; 5 and 10. Table A.4 gives the bias and coverage. Figure A.4 shows the difference in mean inclusion probability between valid and invalid instruments.

**Table A.4:** Evaluation criteria with many weak instruments. 100 instruments in total. True *β* is 0.05. No. inv., Number of invalid instrument(s); *Q*, Q-statistics with exact weights; bias, mean bias; Cover., coverage; DL est., DL estimate; Full Bayes., Full Bayesian; 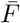, mean F-statistics.

| No. inv. | Q | IVW |  | DL est. |  | Full Bayes. |  | MR-APS |  | MR-RAPS |  |
| --- | --- | --- | --- | --- | --- | --- | --- | --- | --- | --- | --- |
|  |  | Bias | Cover. | Bias | Cover. | Bias | Cover. | Bias | Cover. | Bias | Cover. |
| $\overline{\mathbf{F}} = 10$ | | | | | | | | | | | |
| 0 | 98.0 | -0.019 | 9.30 | -0.001 | 96.40 | 0.000 | 94.90 | -0.000 | 93.70 | 0.000 | 93.10 |
| 20 | 109.3 | -0.019 | 13.80 | -0.001 | 96.30 | 0.003 | 92.90 | 0.003 | 90.10 | 0.002 | 91.50 |
| 40 | 120.2 | -0.019 | 14.10 | -0.000 | 96.70 | 0.006 | 89.50 | 0.007 | 84.10 | 0.006 | 85.90 |
| 60 | 131.0 | -0.019 | 21.20 | 0.000 | 95.90 | 0.008 | 81.40 | 0.009 | 72.00 | 0.009 | 74.20 |
| 80 | 139.5 | -0.019 | 21.10 | 0.000 | 96.40 | 0.012 | 73.20 | 0.012 | 62.10 | 0.011 | 65.10 |
| 100 | 149.4 | -0.019 | 29.10 | 0.002 | 94.10 | 0.016 | 62.40 | 0.016 | 49.70 | 0.016 | 50.00 |
| $\overline{\mathbf{F}} = 5$ | | | | | | | | | | | |
| 0 | 97.8 | -0.038 | 0.00 | -0.006 | 94.30 | 0.001 | 90.40 | 0.000 | 93.20 | 0.000 | 93.10 |
| 20 | 104.8 | -0.039 | 0.00 | -0.005 | 95.40 | 0.004 | 87.80 | 0.003 | 92.40 | 0.003 | 92.30 |
| 40 | 110.7 | -0.038 | 0.00 | -0.004 | 94.60 | 0.009 | 83.00 | 0.009 | 87.40 | 0.008 | 89.30 |
| 60 | 116.8 | -0.038 | 0.00 | -0.001 | 97.60 | 0.012 | 74.20 | 0.013 | 80.00 | 0.013 | 81.80 |
| 80 | 121.2 | -0.038 | 0.00 | 0.001 | 96.30 | 0.016 | 68.90 | 0.015 | 74.50 | 0.016 | 74.40 |
| 100 | 125.7 | -0.038 | 0.00 | 0.003 | 97.10 | 0.021 | 55.20 | 0.022 | 59.30 | 0.022 | 59.90 |

**Fig. A.4:**
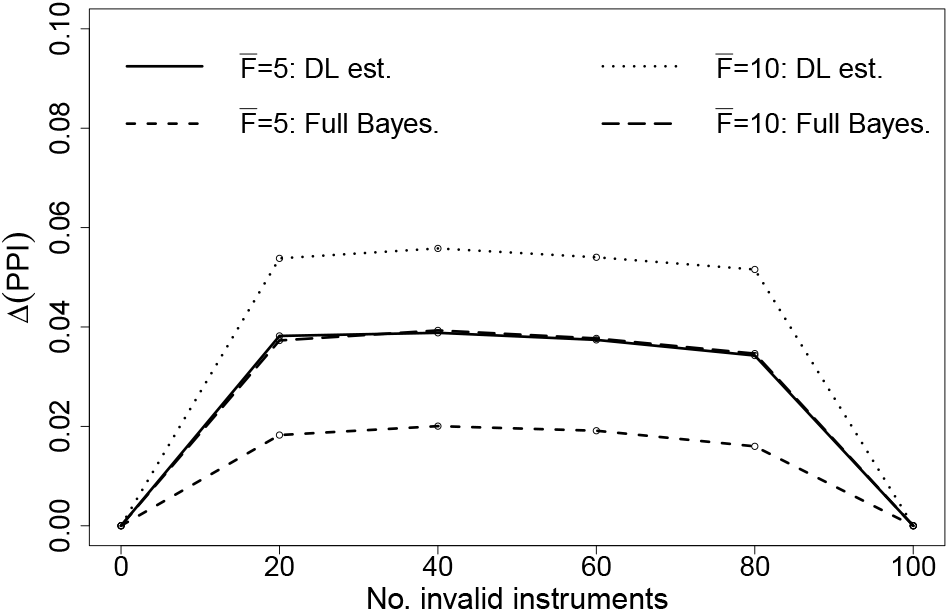
The difference in mean *Δ*(*PPI*) between valid and invalid instruments for many weak instruments. As shown by the legend: solid and short dashed lines are when instruments have mean F-statistics 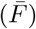 of 5 for DL estimate and full Bayesian respectively. Dotted and long dashed lines are 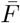 of 10 for DL estimate and full Bayesian respectively.

### E.5 Sensitivity to strengths of heterogeneity

We can use Q-statistics to monitor the heterogeneity between the causal effect estimate from each of the instruments [8]. This section investigates our approaches’ sensitivity to the change in Q-statistics. Using Equation (18) and the *χ*^2^ distribution for *L* − 1 degrees of freedom, we could fix 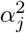 to give p-values for different levels of heterogeneity. We considered 2 forms of Q-statistics; (1) the true Q-statistics in total for 20% invalid instruments are 85, 75, 66 and 62 to give p-value of 0.001, 0.01, 0.05 and 0.1 respectively. (2) Each invalid instruments have true Q-statistics of 11, 7, 4 and 3 to give p-value of 0.001, 0.01, 0.05 and 0.1 respectively. But in total, it is borderline evidence for heterogeneity (Q-statistic p-value=0.05), hence, the number of invalid instruments increases with the Q-statistics. See Table A.5 for a summary.

**Table A.5:** Summary of Q-statistics (*Q*) simulation. The p-value for overall and individual *Q* is from *χ*^2^ distribution of *L*−1 and 1 degrees of freedom respectively. Total number of instruments is 50. Ind., individual.

| Scenario | No. invalid | Overall Q (p-value) | Individual Q (p-value) |
| --- | --- | --- | --- |
| Sum $Q$ | 10 | 85 (0.001) | 8.5 (0.01) |
|  | 10 | 75 (0.01) | 7.5 (0.01) |
|  | 10 | 66 (0.05) | 6.6 (0.01) |
|  | 10 | 62 (0.1) | 6.2 (0.01) |
| Ind. $Q$ | 6 | 66 (0.05) | 11 (0.001) |
|  | 10 | 66 (0.05) | 7 (0.01) |
|  | 17 | 66 (0.05) | 4 (0.05) |
|  | 25 | 66 (0.05) | 3 (0.1) |

Our results demonstrate three facts:

1. Increasing heterogeneity with same number of invalid instruments does not affect the overall performance of the estimators, but only the inclusion probability of the instruments.
2. Increasing the number of invalid instruments whilst fixing the total heterogeneity does not affect the overall performance of the estimators.
3. When the pleiotropy parameters are small and exchangeable, the probability of inclusion is approximately constant across SNPs

**Table A.6:** Evaluation criteria for varying Q-statistics. 50 instruments with mean F-statistics of 100. True *β* is 0.05. No. inv., Number of invalid instrument(s); *Q* exact, estimated Q-statistics for all instruments with exact weights; Cover., coverage; Ind., individual.

| Scenario | No. inv. | Overall Q | Ind. Q | Q exact | DL est. |  | Full Bayes. |  |
| --- | --- | --- | --- | --- | --- | --- | --- | --- |
|  |  |  |  |  | Bias | Cover. | Bias | Cover. |
| Sum $Q$ | 10 | 85 | 8.5 | 134.5 | 0.001 | 94.90 | 0.001 | 92.30 |
|  | 10 | 75 | 7.5 | 124.2 | 0.000 | 97.00 | 0.000 | 94.40 |
|  | 10 | 66 | 6.6 | 115.3 | 0.001 | 96.10 | 0.001 | 93.60 |
|  | 10 | 62 | 6.2 | 111.0 | 0.000 | 96.60 | 0.001 | 94.90 |
| Ind. $Q$ | 6 | 66 | 11 | 113.8 | 0.000 | 96.80 | 0.000 | 95.90 |
|  | 10 | 66 | 7 | 115.4 | 0.001 | 96.60 | 0.001 | 94.00 |
|  | 17 | 66 | 4 | 113.7 | -0.001 | 95.90 | 0.000 | 91.70 |
|  | 25 | 66 | 3 | 116.7 | -0.001 | 96.00 | 0.000 | 87.60 |

**Fig. A.5:**
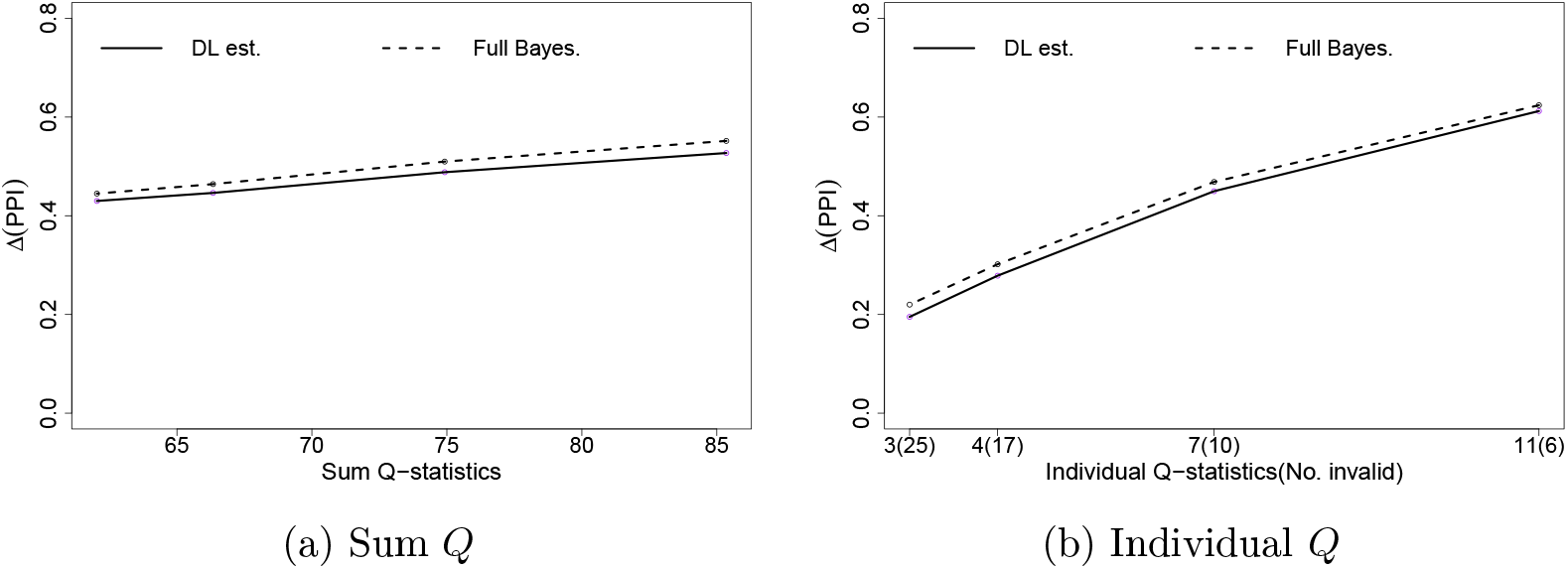
The difference in mean *Δ*(*PPI*) for (a) sum *Q* of all invalid instruments (b) when the fixed amount of heterogeneity (*Q*=66) is due to many weakly pleiotropic or a small number of highly pleiotropic SNPs. As shown by legend: solid and short dashed lines are DL estimate and full Bayesian respectively.

## F Modified Metropolis-Hastings algorithm for InSIDE violating pleiotropy

The updating algorithm for *β*_1_, *β*_2_, 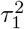 and 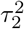 is the same as *β* and *τ*^2^ in the one-parameter model respectively (Appendix C).

– **Update** *I*_1_

1. Generate a random number between 1 and *L*, define it as 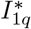 from *P*(*I*_*L*_), which is the *q*th element of 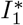
2. Set 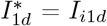 for all *d* ≠ *q*, if *I*_*i*2*q*_ ≠ 1, otherwise repeat step 1.
3. Set 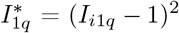.
4. If 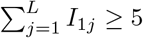 is true, proceed to next step, otherwise repeat step 1.
5. Accept 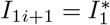 with probability:

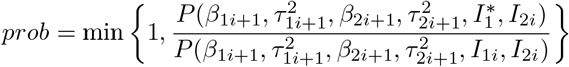

otherwise set *I*_1*i*+1_ = *I*_1*i*_.
– **Update** *I*_2_

1. Generate a random number between 1 and *L*, define it as 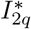 from *P*(*I*_*L*_), which is the *q*th element of 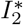
2. Set 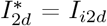 for all *d* ≠ *q*, if *I*_*i*1*q*_ ≠ 1, otherwise repeat step 1.
3. Set 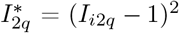.
4. If 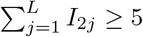 is true, proceed to next step, otherwise repeat step 1.
5. Accept 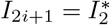 with probability:

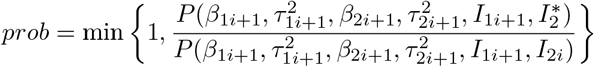

otherwise set *I*_2*i*+1_ = *I*_2*i*_.

Step 2 in **Update** *I*_1_ and **Update** *I*_2_ restricts the new jump to be conditional on *I*_2_ and *I*_1_ respectively, this will stop the case of (*I*_1*j*_ = 1, *I*_2*j*_ = 1). Model space including both (*I*_1*j*_ = 1, *I*_2*j*_ = 1) and (*I*_1*j*_ = 0, *I*_2*j*_ = 0) is equivalent to giving model that consists of outlying instruments higher probability then models where instruments have to be designated to either *I*_1_ or *I*_2_.

## G Simulations under the two-parameter model

This section is for two-parameter BESIDE-MR, that covers Monte Carlo simulation method, results for weaker instruments (L=50 and mean F-statistics of 10), and simulated example to demonstrate when a SNP belongs to *S*_0_ (neither *I*_1_ or *I*_2_ clusters).

### G.1 Simulation Method

Using the same underlying data generating Model 16, suppose that we have two different groups of invalid instruments: in the first group, *S*_1_ we have *Ψ*_*j*_ = 0 for all SNPs and 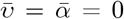, shown in Appendix E. That is, the SNPs in *S*_1_ exhibit balanced pleiotropy under the InSIDE assumption. For illustrative purposes, suppose now that the remaining instruments are in a set *S*_2_, defined by *δ*_*j*_ = 0, *v*_*j*_ = 0 and *κ*_*x*_ = *κ*_*y*_ = 1, but *Ψ*_*j*_ ≠ 0 have Uniform *U*(0.34, 1.1) distribution. This means that that *α*_*j*_ = *γ*_*j*_ = *Ψ*_*j*_, so that the InSIDE assumption is perfectly violated. Using the bias formulae, Equation (2.4) in the main manuscript, it follows that

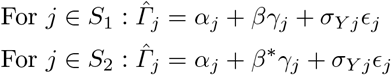

where *β*\* = *β* + 1. The set of SNPs in *S*_2_ therefore identify a distinct, biased version of the causal effect. In the general case where the SNPs could be classified into an InSIDE-respecting set and an InSIDE-violating set, it would be more reasonable to assume that *α*_*j*_, *γ*_*j*_ and *v*_*j*_ could all be non-zero. Although InSIDE would not then be maximally violated in *S*_2_ we would still see two clusters in the data, albeit with a less defined separation.

The same evaluation criteria is used as for the one-parameter model but now *Δ*(*PPI*) is probability of inclusion for *S*_1_ and *S*_2_ SNPs, where their numbers add up to *L*.

### G.2 Weak instruments

We reduced the strength of instrument of scenario 6 to have mean F-statistics of 10; *σ*_*Xj*_ are generated from a Uniform *U*(0.06, 1) distribution for both *S*_1_ and *S*_2_. Table A.7 gives the bias and coverage. Figure A.6 shows the difference in mean probability of inclusion between *S*_1_ and *S*_2_ instruments.

### G.3 Simulated example for S_0_

**Table A.7:** Evaluation criteria for estimating two causal parameter from instruments with mean F-statistic of 10. 50 instruments in total. The true *β* is 0.05. Est., estimator; Inst., instrument(s); *Q*, exact Q-statistics; DL est., DL estimate; Full Bayes., Full Bayesian. *β*_1_ is estimating *β* and *β*_2_ for *β* + 1.

| Est. | Inst.<br>$S_1 : S_2$ | Q | | mean bias | | median bias | | coverage | |
| --- | --- | --- | --- | --- | --- | --- | --- | --- | --- |
| | | $S_1$ | $S_2$ | $\beta_1$ | $\beta_2$ | $\beta_1$ | $\beta_2$ | $\beta_1$ | $\beta_2$ |
| DL est. | 40:10 | 58.8 | 51.2 | -0.004 | -0.988 | -0.003 | -0.990 | 98.7 | 0.2 |
|  | 30:20 | 43.5 | 118.9 | 0.026 | -0.870 | 0.014 | -0.974 | 98.4 | 18.3 |
|  | 25:25 | 35.4 | 153.2 | 0.051 | -0.532 | 0.020 | -0.511 | 95.9 | 60.4 |
|  | 20:30 | 28.0 | 205.5 | 0.027 | -0.252 | 0.012 | -0.185 | 94.8 | 86.8 |
|  | 10:40 | 12.8 | 278.5 | 0.252 | -0.143 | 0.178 | -0.118 | 75.3 | 93.9 |
| Full Bayes. | 40:10 | 58.8 | 51.2 | -0.236 | -0.478 | -0.002 | -0.050 | 81.4 | 70.3 |
|  | 30:20 | 43.5 | 118.9 | -0.395 | -0.364 | -0.009 | 0.064 | 65.9 | 69.8 |
|  | 25:25 | 35.4 | 153.2 | 0.551 | -1.420 | 0.973 | -1.025 | 30.1 | 41.4 |
|  | 20:30 | 28.0 | 205.5 | 0.500 | -1.566 | 1.062 | -1.816 | 19.0 | 42.4 |
|  | 10:40 | 12.8 | 278.5 | 0.258 | -2.332 | 1.087 | -3.133 | 1.9 | 27.2 |

**Fig. A.6:**
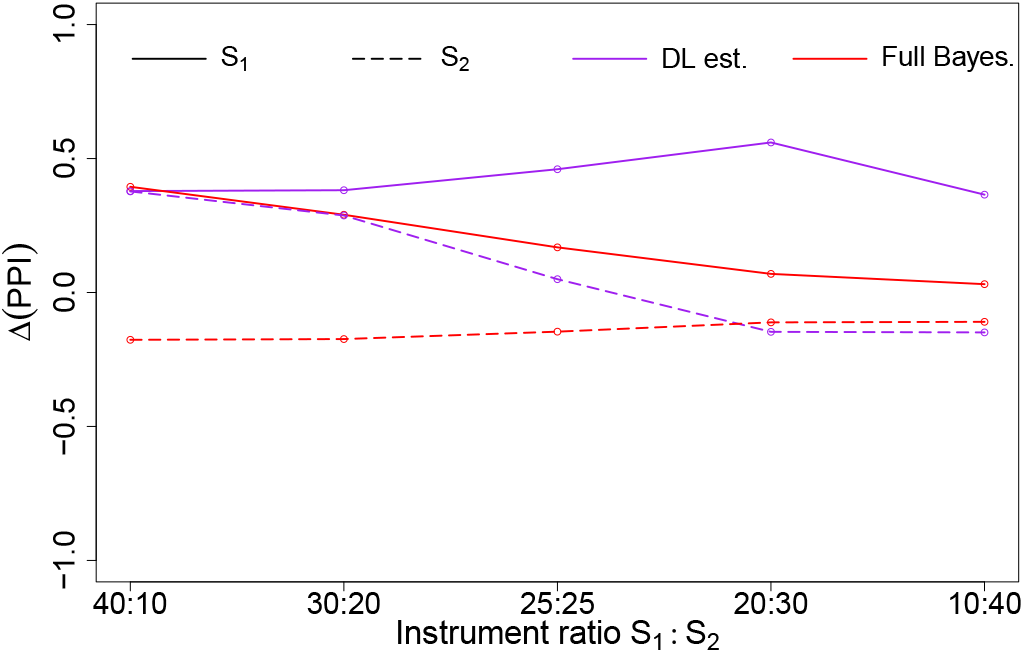
Mean difference in the *Δ*(*PPI*) between *S*_1_ and *S*_2_ as a function of the true ratio *S*_1_:*S*_2_ for weak instruments (mean F-statistic of 10).

**Fig. A.7:**
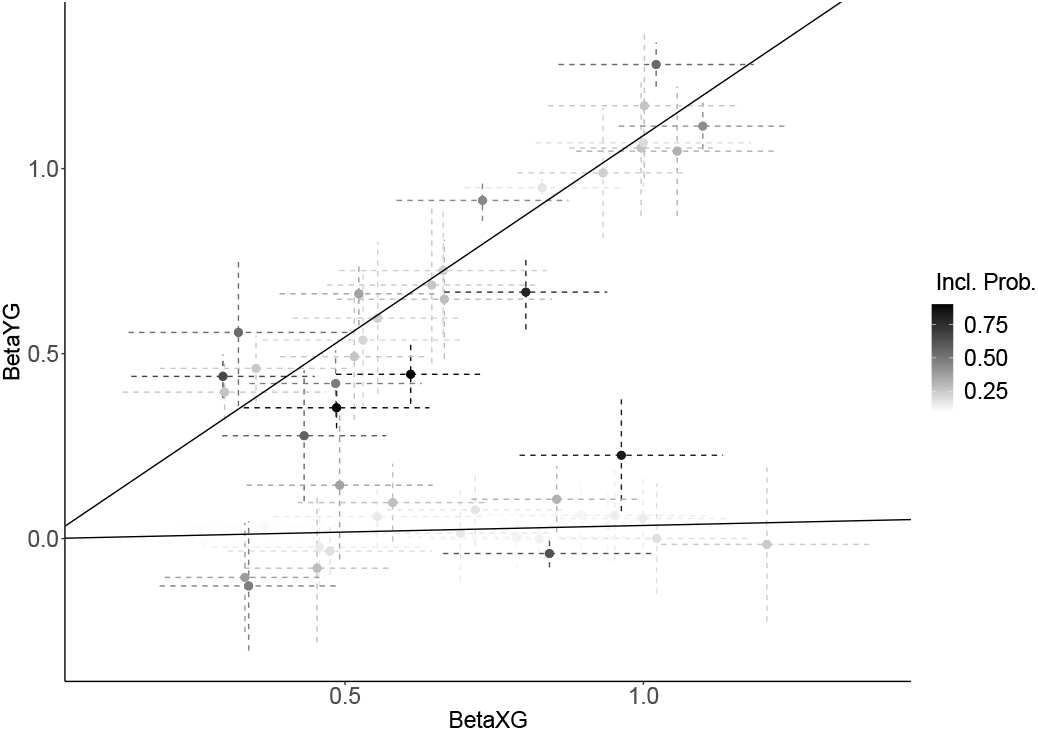
An association plot of a simulated example to demonstrate when a SNP is in *S*_0_ (neither *S*_1_ or *S*_2_). The simulated *S*_1_:*S*_2_ ratio is 50:50 for strong instruments (mean F-statistic of 100). The 2 solid lines are the DL estimated effect sizes for the 2 clusters. As shown by legend; the colour gradient is the *PPI* for a instrument belonging to *S*_0_, i.e. the darker the colour the higher the probability that the SNP belongs to *S*_0_.

## H Applied example

This section gives the *PPI* from two-parameter BESIDE-MR for each SNP. And results from sensitivity analysis for both one- and two-parameter BESIDE-MR.

**Fig. A.8:**
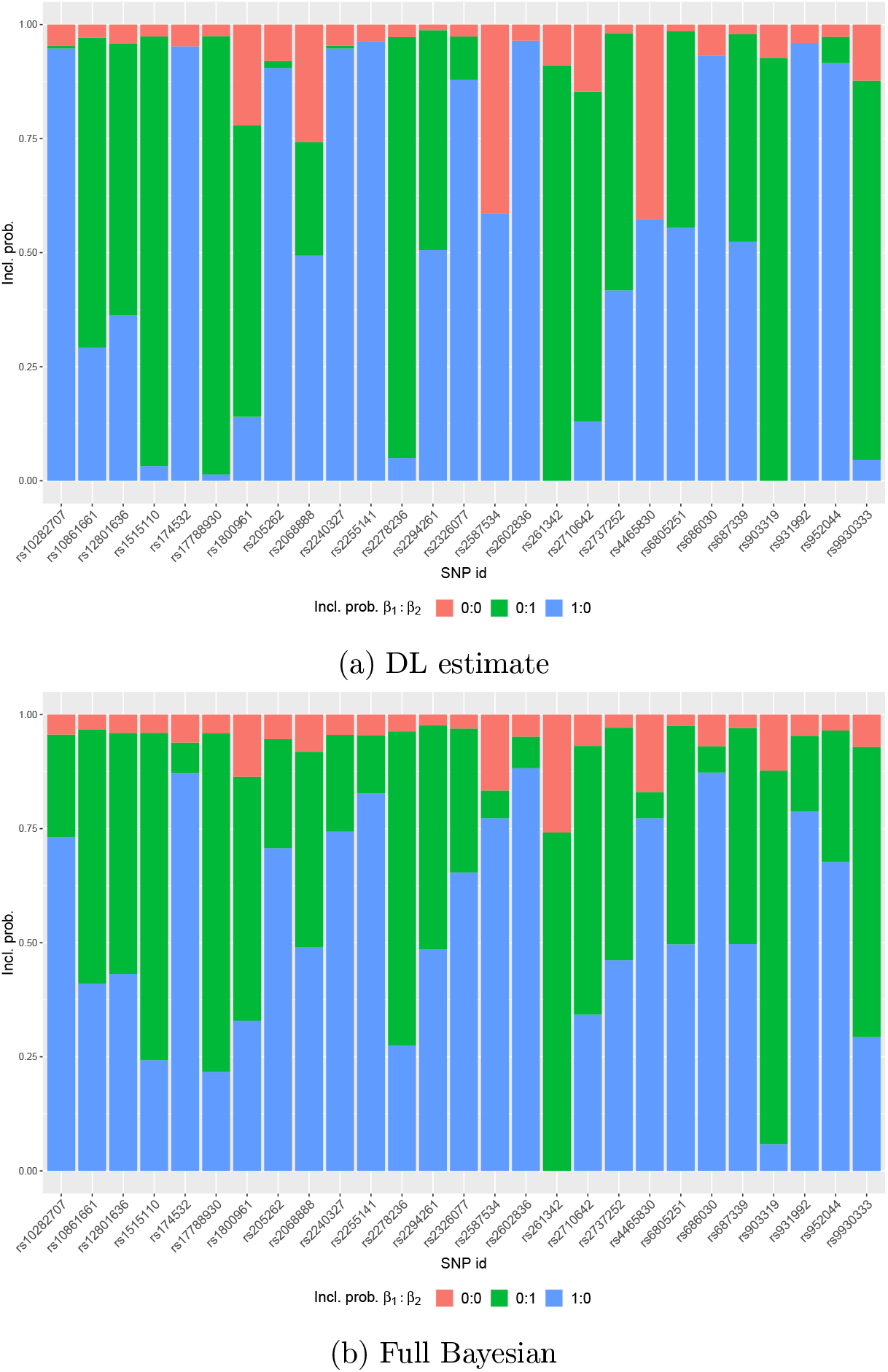
AMD and HDL: *PPI* for DL estimate (a) and full Bayesian approach (b), assuming InSIDE violation. As shown by legend; colour red, green and blue is for instrument in neither (0:0), instrument estimating *β*_2_ (0:1) and *β*_1_ (1:0) respectively.

**Table A.8:** Sensitivity analysis for one-parameter BESIDE-MR with non-zero penalisation term, *η*. Med., LCI and UCI are the median of the posterior distribution with 95% upper and lower credible intervals respectively. 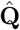 is instrument normalised Q-statistics, Σ**Q**_**j**_/**I**_**j**_. Σ*I*_*j*_ is the number of instruments included. The Q-statistic for 27 Instruments is 115.99.

| $\eta$ | | **2** | | | **3** | | | **4** | | | **5** | | |
| --- | --- | --- | --- | --- | --- | --- | --- | --- | --- | --- | --- | --- | --- |
| **Para.** | **Est.** | **Med.** | **LCI** | **UCI** | **Med.** | **LCI** | **UCI** | **Med.** | **LCI** | **UCI** | **Med.** | **LCI** | **UCI** |
| $\beta$ | DL | 0.78 | 0.53 | 1.08 | 0.79 | 0.53 | 1.08 | 0.78 | 0.53 | 1.07 | 0.78 | 0.53 | 1.07 |
|  | Bayes | 0.77 | 0.50 | 1.08 | 0.77 | 0.50 | 1.07 | 0.77 | 0.50 | 1.07 | 0.76 | 0.49 | 1.07 |
| $\tau^2 \times 10^{-4}$ | DL | 0.00 | 0.00 | 0.00 | 0.00 | 0.00 | 0.00 | 0.00 | 0.00 | 0.00 | 0.00 | 0.00 | 0.00 |
|  | Bayes | 0.28 | 0.09 | 1.77 | 0.29 | 0.09 | 1.83 | 0.29 | 0.09 | 1.86 | 0.29 | 0.09 | 1.87 |
| $\hat{\mathbf{Q}}$ | DL | 0.98 | 0.89 | 0.99 | 0.98 | 0.93 | 0.99 | 0.99 | 0.94 | 0.99 | 0.99 | 0.94 | 0.99 |
|  | Bayes | 1.86 | 1.51 | 1.94 | 1.86 | 1.57 | 1.90 | 1.86 | 1.57 | 1.86 | 1.86 | 1.86 | 1.86 |
| $\sum I_j$ | DL | 20 | 19 | 20 | 20 | 19 | 20 | 20 | 20 | 20 | 20 | 19 | 20 |
|  | Bayes | 26 | 24 | 26 | 26 | 25 | 26 | 26 | 25 | 26 | 26 | 25 | 26 |

**Table A.9:** Sensitivity analysis for two-parameter model with non-zero penalisation terms, *η*_1_ and *η*_2_. Median, 95% LCI and 95% UCI are the median of the posterior distribution with 95% upper and lower credible intervals respectively. 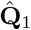 and 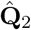 are the instrument normalised Q-statistics, Σ**Q**_**1j**_/**I**_**1j**_ and Σ**Q**_**2j**_/**I**_**2j**_ respectively. Σ*I*_1*j*_ and Σ*I*_2*j*_ are the number of instruments included in the 2 clusters. The Q-statistic for 27 instruments is 115.99.

| $\vert \eta_1 - \eta_2\vert $<br>Para. | Est. | 1 | | | 2 | | | 3 | | | 4 | | | 5 | | |
| --- | --- | --- | --- | --- | --- | --- | --- | --- | --- | --- | --- | --- | --- | --- | --- | --- |
|  |  | Med. | LCI | UCI | Med. | LCI | UCI | Med. | LCI | UCI | Med. | LCI | UCI | Med. | LCI | UCI |
| $\beta_1$ | DL | 0.91 | -1.03 | 1.84 | 0.96 | -0.77 | 1.86 | 0.93 | 0.57 | 1.67 | 0.93 | 0.54 | 1.67 | 0.89 | 0.58 | 1.50 |
|  | Bayes | 0.86 | -0.85 | 1.84 | 0.87 | -0.72 | 1.82 | 0.86 | 0.42 | 1.57 | 0.86 | 0.43 | 1.58 | 0.84 | 0.52 | 1.36 |
| $\beta_2$ | DL | -0.52 | -1.72 | 1.75 | -0.67 | -1.83 | 1.52 | -0.79 | -1.86 | 0.49 | -0.78 | -1.87 | 1.00 | -0.82 | -1.83 | -0.04 |
|  | Bayes | 0.43 | -1.61 | 1.90 | 0.21 | -1.71 | 1.89 | -0.53 | -1.80 | 1.75 | -0.56 | -1.82 | 1.74 | -0.74 | -1.79 | 1.49 |
| $\tau_1^2 \times 10^{-4}$ | DL | 0.00 | 0.00 | 0.00 | 0.00 | 0.00 | 0.00 | 0.00 | 0.00 | 0.00 | 0.00 | 0.00 | 0.00 | 0.00 | 0.00 | 0.00 |
|  | Bayes | 0.24 | 0.08 | 1.30 | 0.24 | 0.08 | 1.26 | 0.24 | 0.08 | 1.24 | 0.24 | 0.08 | 1.26 | 0.24 | 0.08 | 1.29 |
| $\tau_2^2 \times 10^{-4}$ | DL | 0.00 | 0.00 | 0.00 | 0.00 | 0.00 | 0.00 | 0.00 | 0.00 | 0.00 | 0.00 | 0.00 | 0.00 | 0.00 | 0.00 | 0.00 |
|  | Bayes | 0.25 | 0.08 | 1.48 | 0.25 | 0.08 | 1.53 | 0.25 | 0.08 | 1.50 | 0.25 | 0.08 | 1.51 | 0.25 | 0.08 | 1.52 |
| $\hat{\mathbf{Q}}_1$ | DL | 0.82 | 0.33 | 0.99 | 0.82 | 0.35 | 0.99 | 0.90 | 0.57 | 1.00 | 0.90 | 0.57 | 1.00 | 0.94 | 0.72 | 1.00 |
|  | Bayes | 1.26 | 0.51 | 2.15 | 1.25 | 0.53 | 2.11 | 1.36 | 0.82 | 2.01 | 1.37 | 0.84 | 2.02 | 1.44 | 1.03 | 2.01 |
| $\hat{\mathbf{Q}}_2$ | DL | 0.70 | 0.16 | 0.99 | 0.69 | 0.14 | 0.98 | 0.73 | 0.16 | 0.98 | 0.73 | 0.17 | 0.98 | 0.77 | 0.25 | 0.98 |
|  | Bayes | 1.22 | 0.25 | 2.68 | 1.21 | 0.22 | 2.82 | 1.20 | 0.23 | 2.94 | 1.19 | 0.23 | 2.88 | 1.14 | 0.26 | 3.03 |
| $\sum I_{1j}$ | DL | 11 | 6 | 15 | 11 | 7 | 15 | 15 | 11 | 18 | 15 | 11 | 18 | 17 | 14 | 19 |
|  | Bayes | 11 | 7 | 16 | 12 | 7 | 16 | 16 | 12 | 20 | 16 | 12 | 20 | 19 | 16 | 22 |
| $\sum I_{2j}$ | DL | 6 | 5 | 9 | 5 | 5 | 7 | 5 | 5 | 6 | 5 | 5 | 6 | 5 | 5 | 6 |
|  | Bayes | 6 | 5 | 9 | 5 | 5 | 7 | 5 | 5 | 6 | 5 | 5 | 6 | 5 | 5 | 5 |

